# Mature tuft cell phenotypes are sequentially expressed along the intestinal crypt-villus axis following cytokine-induced tuft cell hyperplasia

**DOI:** 10.1101/2024.11.28.625899

**Authors:** Julian R. Buissant des Amorie, Max A. Betjes, Jochem Bernink, Joris H. Hageman, Maria C. Heinz, Ingrid Jordens, Tiba Vinck, Ronja M. Houtekamer, Ingrid Verlaan-Klink, Sascha R. Brunner, Dimitrios Laskaris, Jacco van Rheenen, Martijn Gloerich, Hans Clevers, Jeroen S. van Zon, Sander J. Tans, Hugo J.G. Snippert

**Author notes:** Correspondence should be addressed to H.J.G.S.

## Abstract

Intestinal tuft cells are epithelial sentinels that trigger host defense upon detection of parasite-derived compounds. While representing interesting targets for immunomodulatory therapies in inflammation-driven intestinal diseases, their detailed functioning is poorly understood. Although two distinct intestinal tuft cell types have been described, we reveal common intermediary transcriptomes among tuft cells in mouse and human. Tuft cell-specific reporter knock-ins in organoids show that the two tuft types are sequentially expressed transcriptomic states that represent different maturation stages. Moreover, cytokines interleukin-4 and interleukin-13 only induce lineage specification to Nrep^+^ tuft-1 cells, while BMP and cholinergic signalling advance differentiation towards immune-related ChAT^+^ tuft-2 phenotypes. Functionally, both tuft cell states have chemosensory capacity and respond to stimuli like succinate, but reaction probability increases during tuft cell maturation. Our tuft type-specific reporters and optimized differentiation strategy in organoids provide an experimental platform to study the functioning of tuft cells and their unique chemosensory properties.

## Introduction

Tuft cells are solitary chemosensory epithelial cells that respond to specific environmental stimuli by secretion of effector molecules that initiate appropriate changes in the immune state and physiology of their surroundings. Although they are rare, they attract significant interest especially now that increasing amounts of evidence underscore their crucial role at the interface between the intestinal epithelial barrier and the immune system, making them interesting targets for therapeutic immunomodulation^1–3^. In the intestine, tuft cells detect parasitic helminths and *Tritrichomonas* protists and secrete the cytokine interleukin-25 (IL-25) to initiate the type 2 immune response required for parasitic clearance, characterized by increased mucus production (*weep*) and muscle contractility (*sweep*)^4–6^. More specifically, IL-25 activates innate lymphoid cells (ILC2s) in the lamina propria and triggers the release of type 2 cytokines, including interleukin-13 (IL-13). IL13, together with IL4, acts on epithelial progenitors and biases their differentiation towards goblet and tuft cells, resulting in a tuft-ILC2 feedforward circuit that leads to a strong increase in tuft cell numbers, up to as much as 8% in the case of a helminth infection^4^.

In addition to remodeling of the intestinal epithelium through the tuft-ILC2 circuit, tuft cells also signal directly to nearby epithelial cells. For instance, tuft cells are the only epithelial cells that express choline acetyltransferase (ChAT), the enzyme catalyzing acetylcholine biosynthesis. In turn, tuft cell-derived acetylcholine induces fluid secretion from enterocytes by stimulating apical Cl-ion release, further contributing to the ‘*weep’* into the intestinal lumen^7^. Similarly, prostaglandin D2 secreted by activated tuft cells can increase mucus secretion from goblet cells^8^.

Following the discovery of tuft cells as important sentinels and signal relays, multiple studies have shed light on the mechanisms of tuft lineage specification. For instance, the transcription factor POU class 2 homeobox 3 (POU2F3) was found to be a master regulator of tuft cell fate and other transcription factors, such as Gfi1b and Sox4, were found to be important for tuft lineage commitment^5,9,10^. Interestingly, while loss of Notch signaling followed by expression of atonal bHLH transcription factor 1 (*Atoh1*) is essential for differentiation towards most secretory cell types (goblet, enteroendocrine and Paneth), the role of Notch signaling and *Atoh1* in tuft cell differentiation seems to be more complex, with multiple studies describing both *Atoh1*-dependent and *Atoh1*-independent tuft cell populations^9–13^.

Despite these insights into early tuft cell fate specification, much less is known about the signals that further advance tuft cell differentiation and maturation, nor how functionality relates to transcriptional heterogeneity that was observed within the intestinal tuft cell pool^14^. Indeed, two different types of tuft cell have been identified by single-cell RNA sequencing: tuft-1 cells with a neuron-related transcriptional program and tuft-2 cells with an immune-related transcriptional program. Computational analysis of spatial transcriptional programs along the crypt-villus axis, suggested zonation of these two tuft types with tuft-2 markers expressed higher up towards the tip of the villus and tuft-1 markers towards the bottom of the villus^15^. While a tuft-1 cell specific function in the small intestine has yet to be found, sensing of the bacterial metabolite N-undecanoylglycine through vomeronasal receptor Vmn2r26, has been mainly attributed to tuft-2 cells^8^. Although multiple metabolite GPCRs, such as *Ffar3* and *Tas1rs*, as well as orphan GPCRs, are specifically expressed by intestinal tuft cells^14^, tuft cell activation by ligand stimulation has only been established for the succinate receptor SUCNR1 and the aforementioned vomeronasal receptor Vmn2r26^8,16^.

Here, we generated and compared single-cell intestinal tuft cell transcriptomes from mice and organoids and developed optimized organoid models with tuft cell reporter knock-ins. These data reveal that tuft cells undergo a linear differentiation trajectory where cells mature from a tuft-1 to a tuft-2 transcriptomic state after stimulation with IL-4 and IL-13. Moreover, we demonstrate tuft cell-intrinsic and stromal signaling cues required to continue maturation of stalled lineage committed tuft-1 cells to tuft-2 cells. Generally, we show that organoid models provide tuft cells that closely resemble *in vivo* tuft phenotypes, including for the human intestine, and are amenable for in-depth cell biological studies, such as real-time imaging of tuft cell behavior and chemosensing capabilities.

## Results

### scRNA-seq reveals a continuum of zonated tuft cell states on the crypt-villus axis

In order to characterize intestinal tuft cell heterogeneity with high-resolution, we performed single-cell RNA sequencing on a FACS-enriched tuft cell population isolated from choline acetyltransferase^Bac^ (*ChAT*^*Bac*^)-eGFP mice where tuft cells are labelled by GFP^17^ (Fig. 1a; Extended Data Fig. 1a). Unsupervised clustering revealed three clusters (Fig. 1a), each with clear expression of general tuft cell markers such as *Dclk1, Trpm5, Avil, Chat*, and *Pou2f3*, indicating successful isolation of genuine tuft cells (Fig. 1b). The two largest identified clusters showed high similarity to genetic signatures of previously described tuft-1 and tuft-2 populations^14^ (Fig. 1c). In addition, we observed a third cluster which was characterized by a slightly lower *ChAT* expression (Fig. 1d) and GFP intensity (Extended Data Fig. 1b). Differential expression analysis showed that this third cluster separated from the other two clusters based on gene expression that is usually also detected in other cellular lineages, such as *Aldob* and *Dmbt1* (enterocytes), *Muc3* (Goblet cells) or *Reg3b* and *Reg3*g (Paneth cells), leading to a relatively low tuft-specific signature score (Fig. 1e, Extended Data Fig. 1c and Supplementary table S1). Moreover, components required for MHCII-dependent antigen presentation, such as *Cd74, H2-Ab1* and *H2-Aa*, earlier found to be expressed by intestinal stem cells^18^, were also expressed in this cluster (Fig. 1e). Collectively, these data suggest that this cluster represents early-state tuft precursors (tuft-p), which aligns with strong expression of proliferation markers like *Mki67, Pcna, Top2a* and *Mcm2* (Fig. 1f). Further, the integration of our scRNA-seq dataset with previously reported single-cell transcriptomic datasets of mouse small intestine^14,19,20^ (Fig. 1g) identified cells with high tuft-p signatures scores, validating the tuft-p phenotype as a genuine tuft cell transcriptomic state (Fig. 1h).

**Figure 1:**
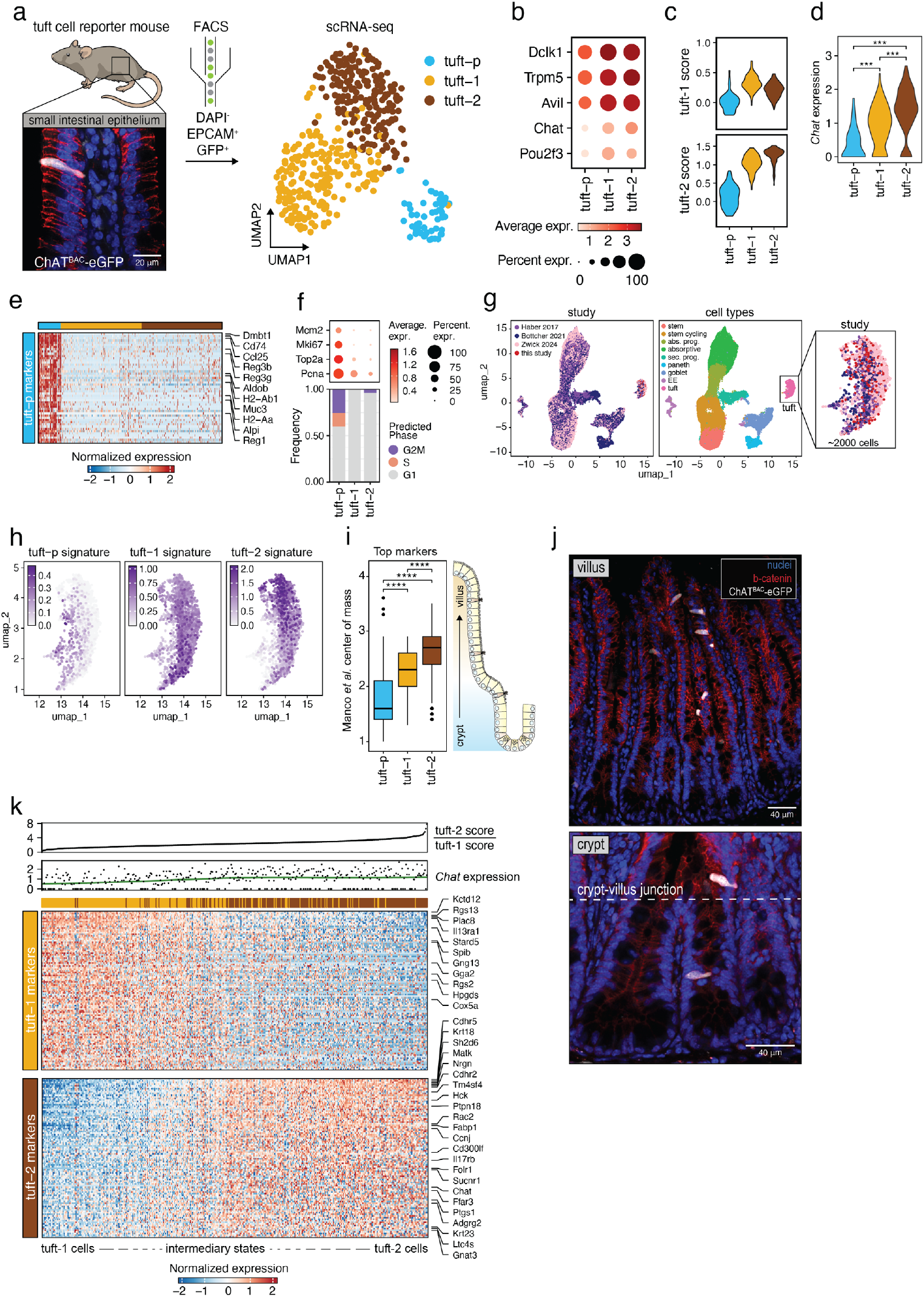
scRNA-seq reveals a continuum of zonated tuft cell states on the crypt-villus axis. **(a)** Left; zoom-in on small intestinal villus where ChAT^BAC^-eGFP marks solitary tuft cells (white), with epithelial membrane (b-catenin, red) and nuclear staining (blue). Right; FACS strategy for plate-based scRNA-seq analysis of GFP^+^ cells from 3 ChAT^BAC^-eGFP mice (pooled) generating UMAP of > 400 ChAT^BAC^-eGFP^+^ tuft cells. Unsupervised clustering resulted in 3 clusters. **(b)** Expression of general tuft cell markers per cluster. Average expression is represented by the dot color while the percentage of expressing cells is denoted by the dot size. **(c)** Violin plots depicting distribution of tuft-1 and tuft-2 signature scores (as described previously^14^) for all cells per cluster. **(d)** Violin plot depicting distribution of relative *Chat* expression for all cells per cluster (**** p < 0.0001, Wilcoxon rank sum test). **(e)** Gene expression profile of tuft-p cells. The top 50 differentially expressed genes (adjusted p-value < 0.01, Wilcoxon rank sum test) with the highest fold-change are plotted and selected genes are highlighted. **(f)** Top; Average expression of proliferation markers (dot color) and percentage of expressing cells (dot size) per cluster. Bottom; distribution of predicted cell-cycle phase per cluster. **(g)** UMAP of integrated single-cell transcriptomic datasets of mouse small intestinal epithelium. Cells are colored by study origin (left) and cell type (middle). Right panel is zoom-in on tuft cell cluster (∼2000 cells). EE: enteroendocrine cells. Abs. prog.: absorptive progenitors. Sec. prog.: secretory progenitors. **(h)** Relative expression of tuft-p, tuft-1 and tuft-2 signatures (top 100 differentially expressed genes) within integrated tuft cell cluster. Signatures as extracted from ChAT-eGFP^+^ tuft clusters in panel a. **(i)** Zonation profile of tuft-p, tuft-1 and tuft-2 clusters from panel a. Boxplots show center of mass on the crypt-villus axis^15^ for the top 100 cluster-specific genes (ANOVA p < 0.0001, **** adjusted p < 0.0001, Tukey HSD test). **(j)** Fluorescent image of ChAT^BAC^-eGFP^+^ (pseudocolor white) mouse small intestine. b-catenin (red) visualizes cell membranes and Hoechst (blue) labels nuclei. Top, representative image showing most ChAT^BAC^-eGFP^+^ tuft cells populate villi. Bottom, representative image showing lower expression level of ChAT^BAC^-eGFP in tuft cells in crypt regions. **(k)** Heatmap showing co-expression of tuft-1 and tuft-2 marker genes (adjusted p-value < 0.01, Wilcoxon rank sum test). Cells from the tuft-1 and tuft-2 clusters from a are plotted and are ordered by the ratio of tuft-1 and 2 scores^14^, indicated at the top of the heatmap. Relative *Chat* expression per cell is shown above the heatmap.

To determine the localization of our three tuft clusters in an unbiased manner, we selected the top marker genes of each cluster and plotted the center of mass of their expression on the crypt-villus axis, as determined by Manco *et al*.^15^ (Fig. 1i). Additionally, we calculated single-cell signature scores along the zones of the crypt-villus axis, to further support regional abundance predictions for the three tuft types^15^ (Extended Data Fig. 1d). This resulted in a zonated pattern, with the tuft-p cells predicted to be closest to the crypt (Fig. 1i and Extended Data Fig. 1d), an intermediate pattern for tuft-1 cells, and the tuft-2 cells located highest up towards the villi tips. Indeed, while *ChAT*^*BAC*^*-*eGFP cells are found mostly on the villi, occasionally *ChAT*^+^ cells are observed in crypts, frequently presenting a dimmer eGFP signal associated with tuft-p cells (Fig. 1j).

Next, taking into account the zonated localization of tuft-1 and tuft-2 cells on the crypt-villus axis, we ranked cells from these clusters based on a tuft-2/tuft-1 signature score and visualized expression patterns of tuft-1 and tuft-2 marker genes in a heatmap (Fig. 1k). Notably, this revealed a gradual transition of tuft-1 to tuft-2 transcriptomic profiles via intermediary phenotypes encompassing mixed expression of both tuft-1 and tuft-2 markers. Crucially, this continuum of cell states and absence of binary confinement, suggests that the two previously reported tuft-types represent sequential differentiation states along the crypt-villus axes. As expected, *Chat* expression levels gradually increased from tuft-1 to tuft-2 states (Fig. 1k), in line with the predominant localization of ChAT^BAC^-GFP^+^ cells towards the higher part of the villus (Fig. 1j). In sum, these descriptive *in vivo* data reveal early tuft precursors located near the crypt and a continuum of tuft-1 and tuft-2 transcriptomic states without binary confinement along the crypt-villus axis.

### Mature small intestinal tuft phenotypes are conserved between mouse and human

To investigate if similar mature tuft cell phenotypes are found in human, we reanalyzed a recently published single-cell transcriptomic dataset of the human small intestinal epithelium, containing ∼800 tuft cells^20^(Fig. 2a). Unsupervised clustering resulted in two tuft clusters, which we labeled tuft-1 and tuft-2 (Fig. 2b) based on differential expression of the mouse tuft-1 and tuft-2 signatures (Fig. 2c). Next, we determined human tuft-1 and tuft-2 signatures and performed GO-term enrichment analysis (Fig. 2d). Akin to mouse, this showed immune-related programs to be enriched in human tuft-2 cells, while human tuft-1 cells showed enrichment of neuron-related programs. Also in the human setting, intermediary tuft phenotypes with cells expressing both tuft-1 and tuft-2 markers could be observed (Fig. 2e). Moreover, previous spatial transcriptomics data^21^ indicate that human tuft cells show a similar zonation profile along the crypt-villus axis as observed in mice, with tuft-2-specific markers found predominantly higher on the crypt-villus axis than tuft-1-specific markers (Fig. 2f).

**Figure 2:**
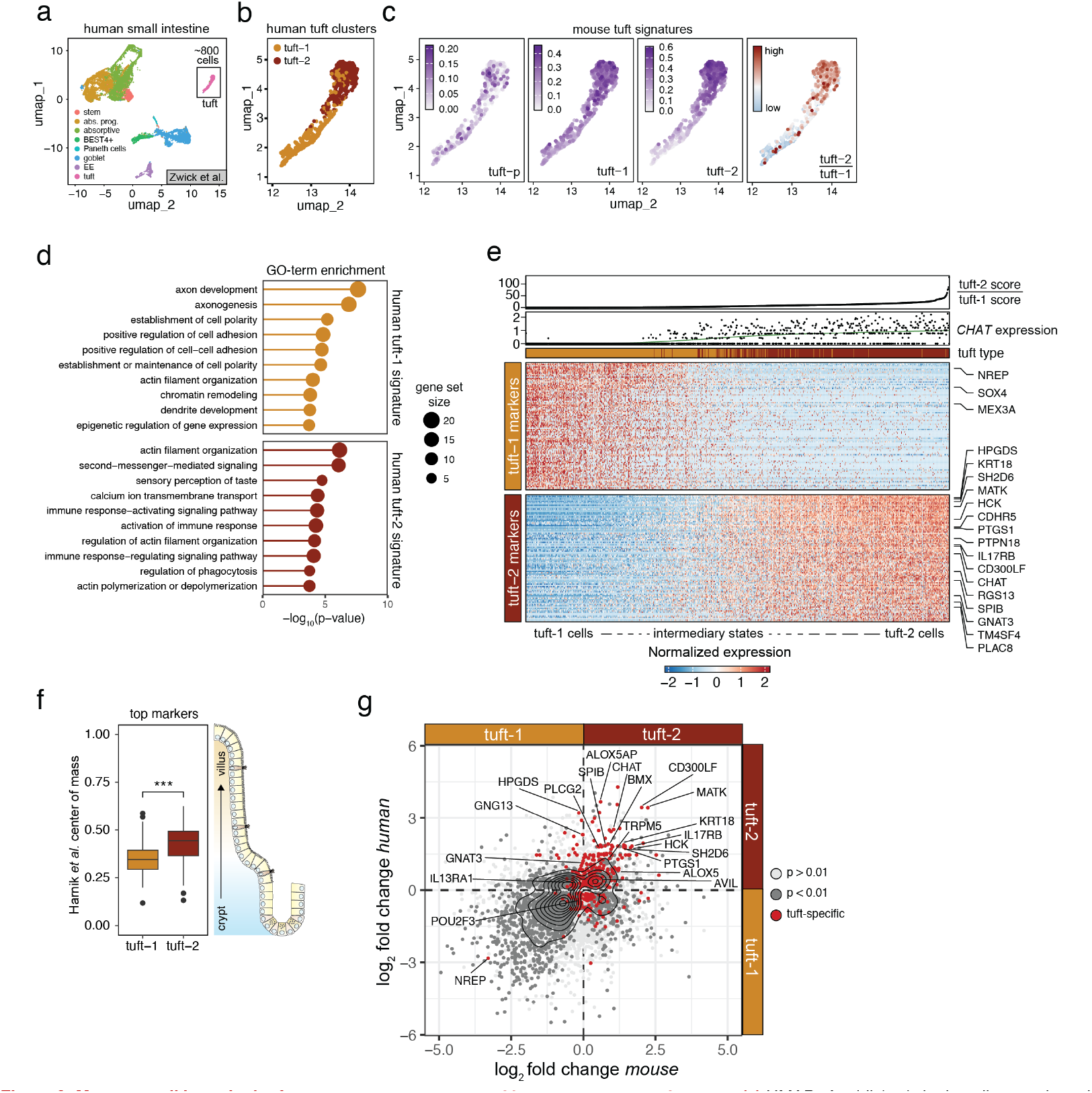
Mature small intestinal tuft phenotypes are conserved between mouse and human. **(a)** UMAP of published single-cell transcriptomics dataset of human small intestine^20^. Cell types are annotated by color. EE: enteroendocrine cells. Abs. prog.: absorptive progenitors. Sec. prog.: secretory progenitors. **(b)** Unsupervised clustering of human tuft cells^20^ identifies two separate tuft cell populations. **(c)** Expression of mouse tuft-p, tuft-1 and tuft-2 signatures extracted from our mouse tuft cell dataset (Fig. 1a). Ratio of mouse tuft-2/tuft-1 scores alligns with the unsupervised clustering of the human tuft cells in panel b. **(d)** GO-term enrichment analysis (clusterProfiler; Biological Process) of human tuft-1 and tuft-2 expression profiles. Dot size represents the number of genes within the gene set. **(e)** Heatmap showing co-expression of tuft-1 and tuft-2 marker genes (adjusted p-value 4 < 0.01, Wilcoxon rank sum test) in human tuft cells. Transcriptomic profiles are ordered by their ratio of human tuft-1 and 2 signature scores, indicated at the top of the heatmap. *CHAT* expression per cell is shown above the heatmap. **(f)** Zonation profile of human tuft-1 and tuft-2 populations from panel b. Center of mass on crypt-villus axis^21^ for the top 50 cluster-specific genes is shown (*** p < 0.001, t-test). **(g)** Differential expression analysis between tuft-1 and tuft-2 clusters in mouse and human. Point color represents significance level (gray). Tones of gray represent significance level, contour plot shows density of significant genes. Tuft-specific genes, in both human and mice, are highlighted in red.

In order to perform a more comprehensive cross-species comparison of human and murine tuft cells, we determined core tuft expression profiles for both species and determined differentially expressed genes between the tuft-1 and tuft-2 clusters (Fig. 2g). Most tuft cell-specific genes were expressed in tuft-2 cells in both mouse and human, suggesting that the tuft-2 state is stronger conserved between mouse and human. Vice versa, most genes that show higher expression levels in tuft-1 cells are not unique to the tuft lineage. Taken together, these analyses show similar transcriptome profiles for mature tuft cells in mice and human and suggest a conserved tuft cell differentiation trajectory in which cells transition from an early tuft-1 phenotype into a mature tuft-2 phenotype.

### Tuft-type specific reporter knock-ins enable identification of tuft-1 and tuft-2 cell states *in vitro*

To experimentally test the differentiation trajectories that give rise to the continuum of tuft-1 and tuft-2 states observed *in vivo* and evaluate their chemosensory capacity, we designed tuft type-specific reporter organoids. This *in vitro* system enables functional investigation of rare tuft cells in a controlled environment and allows for real-time imaging of their differentiation trajectories. For this, we first established small intestinal organoids from *ChAT*^BAC^-eGFP mice, in which spontaneous differentiation towards GFP^+^ tuft cells was occasionally observed in the standard culture medium (Fig. 3a). Notably, unlike DCLK1^+^ tuft cells, the fraction of GFP^+^ tuft cells barely increased in organoids after a commonly applied treatment with IL-4 and IL-13^4,5,22,23^ (Fig. 3bcd). Like *in vivo*, GFP^+^ tuft cells in organoids recapitulated preferential localization to the villus compartment (Fig. 3d). Similar to GFP^+^ cells in *ChAT*^BAC^-eGFP mice, GFP^+^ cells in organoids co-expressed mature tuft-2 markers, such as CD45 and FOLR1 (Extended Data Fig. 2a). In sum, these data suggest that *ChAT*^BAC^-eGFP in organoids, like *in vivo*, preferentially marks a subset of tuft cells that is enriched for mature tuft-2 markers.

**Figure 3:**
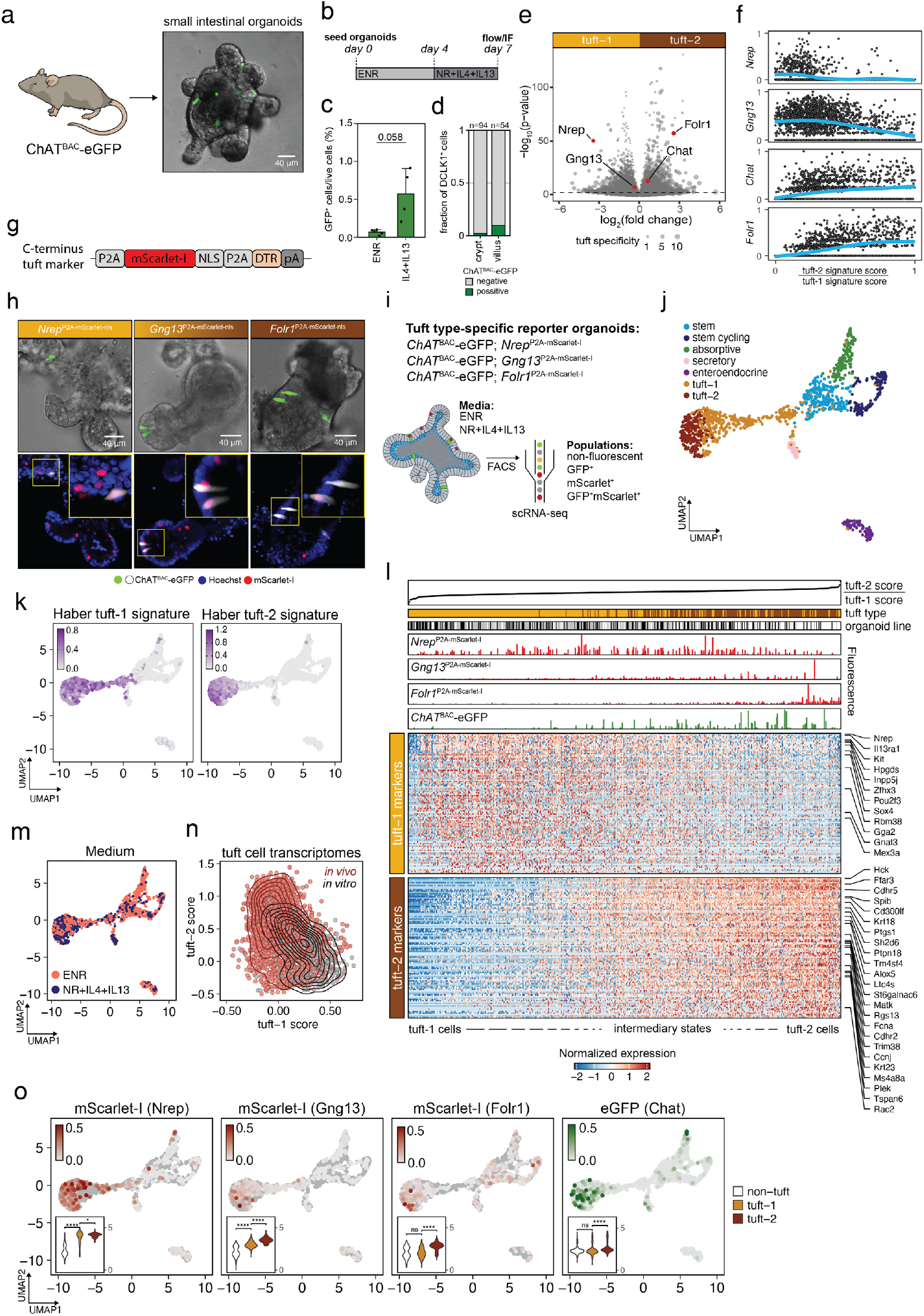
Tuft type-specific reporter knock-ins enable identification of tuft-1 and tuft-2 transcriptional states *in vitro*. **(a)** Small intestinal organoids were derived from ChAT^BAC^-eGFP mice. Phase contrast image of organoid with fluorescent ChAT^BAC^-eGFP^+^ cells in green. **(b)** Treatment regimen for tuft cell induction in organoids with IL-4 and IL13. E, N, R: EGF, R-spondin, Noggin resp. **(c)** Flow cytometry analysis of percentage of GFP^+^ cells in organoids cultured with or without IL-4 and IL-13 according to the regimen shown in b. E, N, R: EGF, R-spondin, Noggin resp. Data are represented as mean ± SD (p-value is shown, t-test). **(d)** Quantification of overlap between GFP^+^ (ChAT^BAC^-eGFP) and DCLK1^+^ cells in organoids as assesed by immunofluorescence following the treatment regimen indicated in b. **(e)** Vulcano plot representing differentially expressed genes (Wilcoxon rank sum test) between tuft-1 and tuft-2 clusters from the integrated mouse *in vivo* single-cell transcriptomic dataset (shown in Fig. 1g). Genes selected for reporter knock-ins are highlighted in red and labeled. Point size represents tuft specificity (see Methods). **(f)** Relative expression levels of the selected reporter genes in all tuft cells from the integrated mouse *in vivo* single-cell transcriptomics dataset (shown in Fig. 1g). Cells are ranked based on their tuft-1/2 signature score ratio to indicate gene expression patterns within tuft population. **(g)** Schematic representation of the knock-in template used to integrate at C-terminus of reporter genes. NLS: nuclear localization signal. mScarlet-I: monomeric red fluorescent protein. DTR: diphtheria toxin receptor. pA: polyadenylation signal. **(h)** Top: Confocal brightfield images of tuft-type-specific reporter knock-in organoids, overlayed with fluorescent ChAT^BAC^-eGFP^+^ tuft cells (green). Bottom: fluorescent overlays of ChAT^BAC^-eGFP^+^ tuft cells (white) with indicated tuft-type reporters (red) and nuclear stain (Hoechst, blue). **(i)** Experimental setup of reporter lines, culture conditions and flow cytometry to enrich for tuft cells prior to single-cell transcriptomic analysis. Indicated fluorescent cell populations were sorted into 384-well plates for plate-based scRNA-sequencing. Fluorescent cells were purposefully overrepresented to enrich for tuft cells. **(j)** UMAP of organoid single-cell transcriptomic dataset, containing cells from all three knock-in tuft reporter lines and both medium conditions. Cell clusters are annotated. **(k)** UMAP of panel h, overlayed with relative tuft-1 and tuft-2 signature scores^14^. **(l)** Heatmap showing co-expression of tuft-1 and tuft-2 marker genes (adjusted p-value < 0.01, Wilcoxon rank sum test) in organoid tuft cells. Organoid tuft cell transcriptomic profiles are ordered by the ratio of mouse tuft-type signature scores, indicated at the top of the heatmap. For each cell the tuft type cluster (yellow: tuft-1; brown: tuft-2) and the organoid knock-in line that it originated from are annotated at the top of the heatmap (black: ChAT^BAC^-eGFP;*Nrep*^P2A-mScarlet-I^, gray: ChAT^BAC^-eGFP;*Gng13*^P2A-mScarlet-I^, white: ChAT^BAC^-eGFP;*Folr1*^P2A-mScarlet-I^). Fluorescent mScarlet-I and eGFP signals of cells during sorting are indicated for each cell at the top of the heatmap (Fluorescence). **(m)** UMAP of organoid single-cell transcriptomic dataset, colored by medium condition. **(n)** Contour plot showing the density of ratios between tuft-1 and tuft-2 signature scores^14^ for tuft cells within the integrated mouse *in vivo* scRNA-sequencing dataset from Fig. 1g (red), as well as the organoids scRNA-seq dataset from Fig. 3j (black), indicating underrepresentation of mature tuft-2 cells *in vitro*. **(o)** mScarlet-I and eGFP fluorescent signal intensities as measured during cell sorting, superimposed on UMAP of organoid single-cell transcriptomic dataset. Insets show the overall fluorescent signal of the reporter per indicated cell cluster (ANOVA p < 0.001, ns not significant, **** adjusted p-value < 0.0001, * adjusted p-value < 0.05, Tukey HSD test).

Next, we used our integrated *in vivo* single-cell transcriptomic datasets of mouse small intestinal epithelium to identify markers that can discriminate between tuft-1 and tuft-2 cells and are not expressed in other epithelial lineages (Fig. 3e and Extended Data Fig. 3ab). We selected *Nrep* as a marker gene that demarcates tuft-1 transcriptomic states (Fig. 3e and Fig. 3f). Vice versa, we picked *Folr1*, which shows highest expression in mature tuft-2 states (Fig. 3e and Fig. 3f). In addition, we chose *Gng13* as an exclusive tuft cell marker, which shows strong expression in tuft-1 and intermediary phenotypes (Fig. 3e and Fig. 3f). Independent knock-ins were integrated at the STOP-codon of these genes and contained a fluorescent mScarlet-I reporter (monomeric RFP variant) and diphtheria toxin receptor (DTR) that both become separated from the tuft cell markers during translation of the P2A peptides^24^ (Fig. 3g and Extended Data Fig. 3cd). Furthermore, we generated all three reporter lines in the *ChAT*^BAC^-eGFP background, such that the eGFP signal can be leveraged as a reference to cross-compare the new tuft-type markers (Fig. 3h).

A prerequisite to study mature tuft cell functioning in organoids, is to confirm that tuft cell phenotypes are accurately recapitulated. To examine the degree of similarity between *in vivo* and *in vitro-*derived tuft cells, we sorted the various fluorescent cell populations from our organoids for scRNA-seq (Fig. 3i and Extended Data Fig. 3e). The ensuing dataset contained >500 tuft cells (Fig. 3j) with tuft-1 and tuft-2 phenotypes that closely resemble their *in vivo* counterparts as illustrated by strong enrichment of tuft-1 and tuft-2 signatures, respectively (Fig. 3k). Resemblance between *in vivo* and *in vitro-*derived tuft cells is further demonstrated by the many tuft cells that showed an intermediary phenotype with mixed expression of both tuft-1 and tuft-2 markers (Fig. 3l). Cytokine treatment did not induce alternative tuft phenotypes that were not present in the standard culture medium, as illustrated by mixing of tuft cells derived from either medium on the UMAP and a lack of medium-specific tuft clusters (Fig. 3m). For a direct comparison, we plotted tuft-1 and tuft-2 signature scores from tuft cells originating from either the integrated *in vivo* dataset (Fig. 1g) or our *in vitro* organoid dataset (Fig. 3j). Despite a strong overlap, most *in vitro* tuft cells were representative of the tuft-1 phenotype (Fig. 3n), presumably a consequence of the crypt-inspired signaling factors in the organoid medium.

Next, we evaluated if our tuft cell type-specific reporters reliably indicate the intended tuft cell transcriptomic states. To this end, we superimposed the signal intensities of the fluorescent reporters, measured during cell sorting, on the single-cell transcriptomics data (Fig. 3o). GFP signal from the *ChAT*^BAC^-eGFP was most dominant in cells with a high tuft-2 signature score (Fig. 3o and Fig. 3l). In contrast, for the tuft-1 marker *Nrep* we found high mScarlet-I signals in tuft-1 cells, which extended to cells with intermediary tuft phenotypes (Fig. 3o and Fig. 3l). As expected, *Gng13* shows a gradual increase of mScarlet-I signal starting with low levels in tuft-1 cells and becoming more abundant towards intermediary and tuft-2 expression profiles (Fig. 3o and Fig. 3l). *Folr1* marks cells with the highest tuft-2 signature scores (Fig. 3o and Fig. 3l). However, despite being predicted as tuft-specific by the *in vivo* single-cell atlases it turned out to be less exclusive for the tuft lineage (Fig. 3o). Nonetheless, *Folr1*-driven mScarlet-I expression is a reliable indicator of tuft-2 phenotypes within the tuft cell compartment and can therefore be used in combination with ChAT to identify the most mature tuft cells (Fig. 3l). Together, these data show that tuft cell phenotypes, including intermediary states, are accurately recapitulated in organoids and they can be readily identified by our tuft type-specific reporters.

### Cytokines drive a unidirectional differentiation trajectory from tuft-1 to tuft-2

To study type 2 cytokine-induced tuft cell differentiation we characterized the frequency of tuft phenotypes in each of the three reporter lines under standard culture conditions (ENR) and upon tuft cell induction with the cytokines IL-4 and IL-13 (IL4+IL13) (Fig. 4a and Fig. 4b). Surprisingly, cytokine treatment predominantly induced a strong increase in the number of mScarlet-I^+^ tuft cells in the *Nrep* and *Gng13* reporter lines, which vastly outnumbered the frequency of ChAT-eGFP^+^ cells (>50 fold) (Fig. 4a and Fig. 4b). On the other hand, the frequency of mScarlet-I^+^ tuft-2 cells in the *Folr1* reporter line increased only mildly and was more reminiscent to the subtle increase in ChAT-eGFP^+^ cells (Fig. 4a and Fig. 4b). These observations are in stark contrast to type 2 cytokine-induced tuft hyperplasia *in vivo*, where both tuft-1 and tuft-2 populations increase in numbers^14,23^ (Extended Data Fig. 4ab). Importantly, this reveals that cytokine administration to organoids, a commonly applied strategy to boost tuft cell numbers, hardly generates tuft-2 cells.

**Figure 4:**
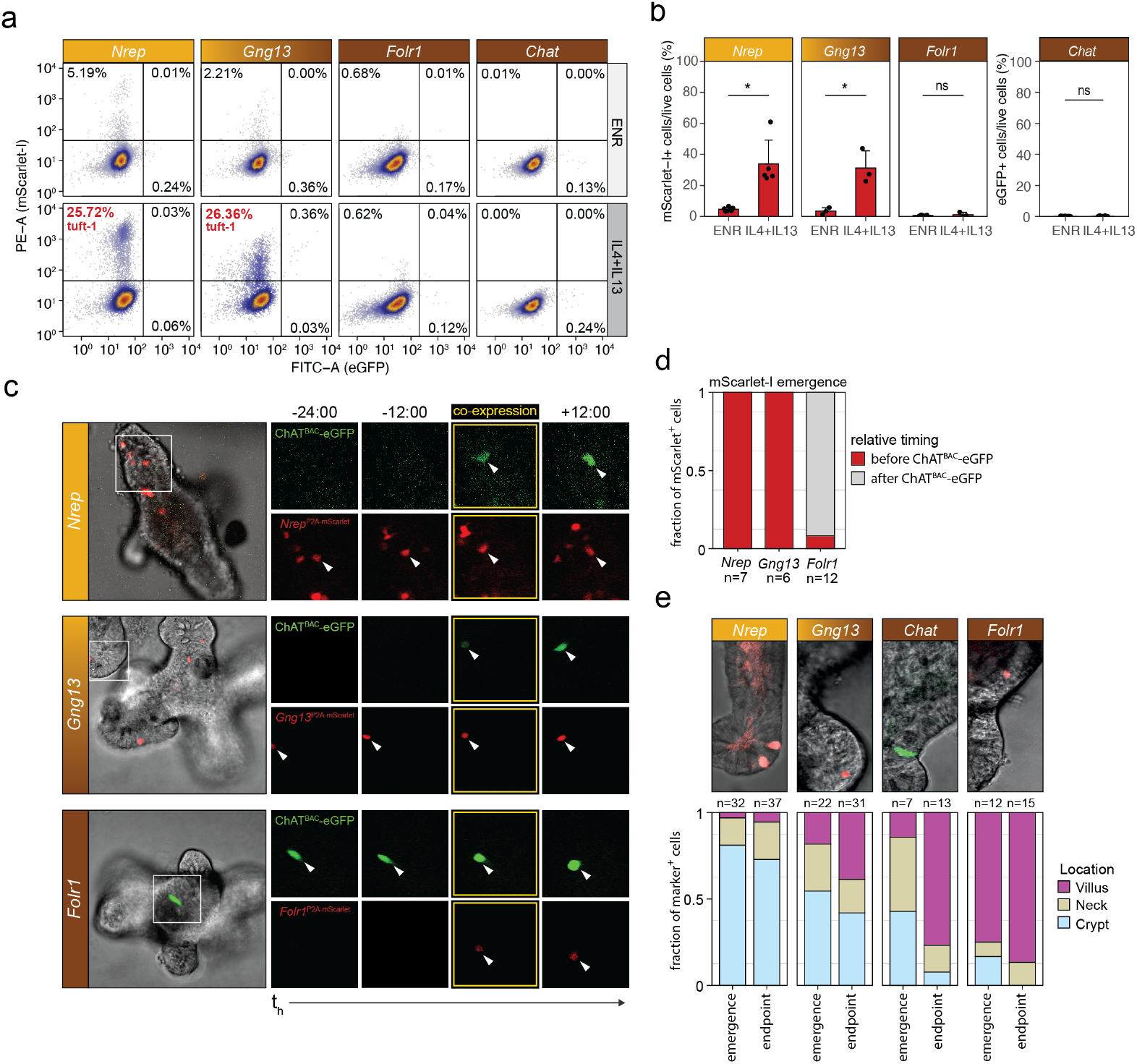
Cytokines drive a unidirectional differentiation trajectory from tuft-1 to tuft-2. **(a)** Representative flow cytometry analysis of the three indicated tuft-type specific reporter lines and the parental ChAT^BAC^-eGFP line (right panel). All lines were treated with IL-4 and IL-13 for 4 days, 3 days after seeding in ENR medium. Percentages of the fluorescent populations are indicated in each plot. Individual panels contain combined data from 3 independent experiments. **(b)** Tuft-type reporter frequencies (%) upon induction with indicated cytokines as in panel a. Flow cytometry quantification are represented as mean ± SD (ns not significant, * p-value < 0.05, t-test). **(c)** Microscopic stills from live imaging experiments demonstrating the temporal dynamics of tuft-type reporters. Left: confocal brightfield image of organoids at the start of imaging, overlayed with fluorescent signals of indicated reporters. Right: stills of fluorescence channels (mSarlet-I: red, eGFP: green) from timepoints preceding and following co-expression of fluorescent markers to visualize differential dynamics of indicated tuft reporters with respect to ChAT^BAC^-eGFP. Cell that shows co-expression is indicated with a white arrowhead throughout imaging timepoints. **(d)** Quantification of mScarlet-I emergence relative to ChAT-eGFP expression for the three indicated tuft type-specific organoid reporter knockins. **(e)** Top: Representative confocal brightfield image with overlay of indicated fluorescent reporter. Bottom: Location distribution of tuft-reporter^+^ cells along the crypt-villus axes within organoids scored at emergence of reporter^+^ cells versus the end of imaging procedure.

Since both the *in vivo* and *in vitro* single-cell transcriptome datasets indicate existence of intermediary tuft phenotypes, we set out to film tuft cell differentiation in our tuft reporter organoids. Generally, we treated organoids with cytokines to induce tuft cell specification and tracked the dynamics of tuft cell identity by means of fluorescent reporters, using confocal live-cell imaging (Fig. 4c). First, we noticed clear differences in temporal expression patterns for the tuft-1 and tuft-2 markers within organoids. Using the tuft marker ChAT^BAC^-eGFP as a reference signal, we observed that tuft-1 marker *Nrep* and tuft1/2 marker *Gng13* become active ahead of *Chat* expression, while the tuft-2 marker *Folr1* becomes active in cells that were already positive for ChAT^BAC^-eGFP (Fig. 4c and Fig. 4d). In agreement with the lack of binary confinement between tuft-1 and tuft-2 cell phenotypes, these temporal expression patterns of tuft-type markers are indicative of a linear differentiation trajectory where tuft-1 cells transition into tuft-2 cells. This is further supported by the fact that tuft-1 marker expression predominantly emerged in the crypt compartment of organoids, while tuft-2 markers appeared in the villus compartment (Fig. 4e). Noticeably, cells positive for our tuft cell markers never underwent mitosis, confirming that even *Nrep*^*+*^ tuft-1 cells are already fully committed to the post-mitotic cell lineage (data not shown). In conclusion, our imaging data reveal that, following IL-4 and IL-13 treatment, tuft-1 cells can transition into tuft-2 cells while migrating to the villus compartment, with no evidence for the opposite trajectory.

### Crypt-villus signaling gradients promote transitioning from tuft-1 to tuft-2 states

Since tuft cell differentiation in cytokine-exposed organoids (IL-4 and IL-13) predominantly resulted in induction of tuft-1 phenotypes (*Nrep* and *Gng13* Fig. 4b) but not tuft-2 phenotypes (*Chat* and *Folr1* Fig. 4b), we set out to identify signals that stimulate the transitioning of tuft-1 cells into mature tuft-2 cells. *In vivo*, tuft-2 markers are found predominantly higher up the crypt-villus axis, which is reflected in our reporter organoids by a higher incidence of *ChAT*^+^ and *Folr1*^+^ tuft-2 cells in the villus compartment of organoids (>50 cells tracked). Therefore, we adjusted the cytokine differentiation medium to better represent a villus environment, starting with the depletion of stem cell-niche inspired growth factors *Noggin, R-spondin* and *EGF* (Fig. 5a). Next, we screened for additional factors that are more abundant in the villus environment, such as *WNT5a, NRG1* and *BMP4*, and factors that are produced by tuft cells and could potentially act in an autocrine manner, such as *BMP2*, acetylcholine and *IL25* (Fig. 5a). Subtraction of the stem cell niche factors (ENR) from the medium resulted in higher frequencies of ChAT^BAC^-eGFP^+^ cells (Fig. 5b). Moreover, both BMPs and acetylcholine further increased the ChAT^BAC^-eGFP^+^ cell population (Fig. 5b). Indeed, according to the single-cell transcriptome datasets, BMP receptors are expressed by tuft cells and the BMP response genes *Id1* and *Id3* are higher expressed in tuft-2 cells (Extended Data Fig. 5a). Concordantly, in mice only the tuft-2, but not the tuft-1 transcriptomic signature, is significantly reduced in *Bmpr1a* depleted intestinal epithelia^25^ (Extended Data Fig. 5b).

**Figure 5:**
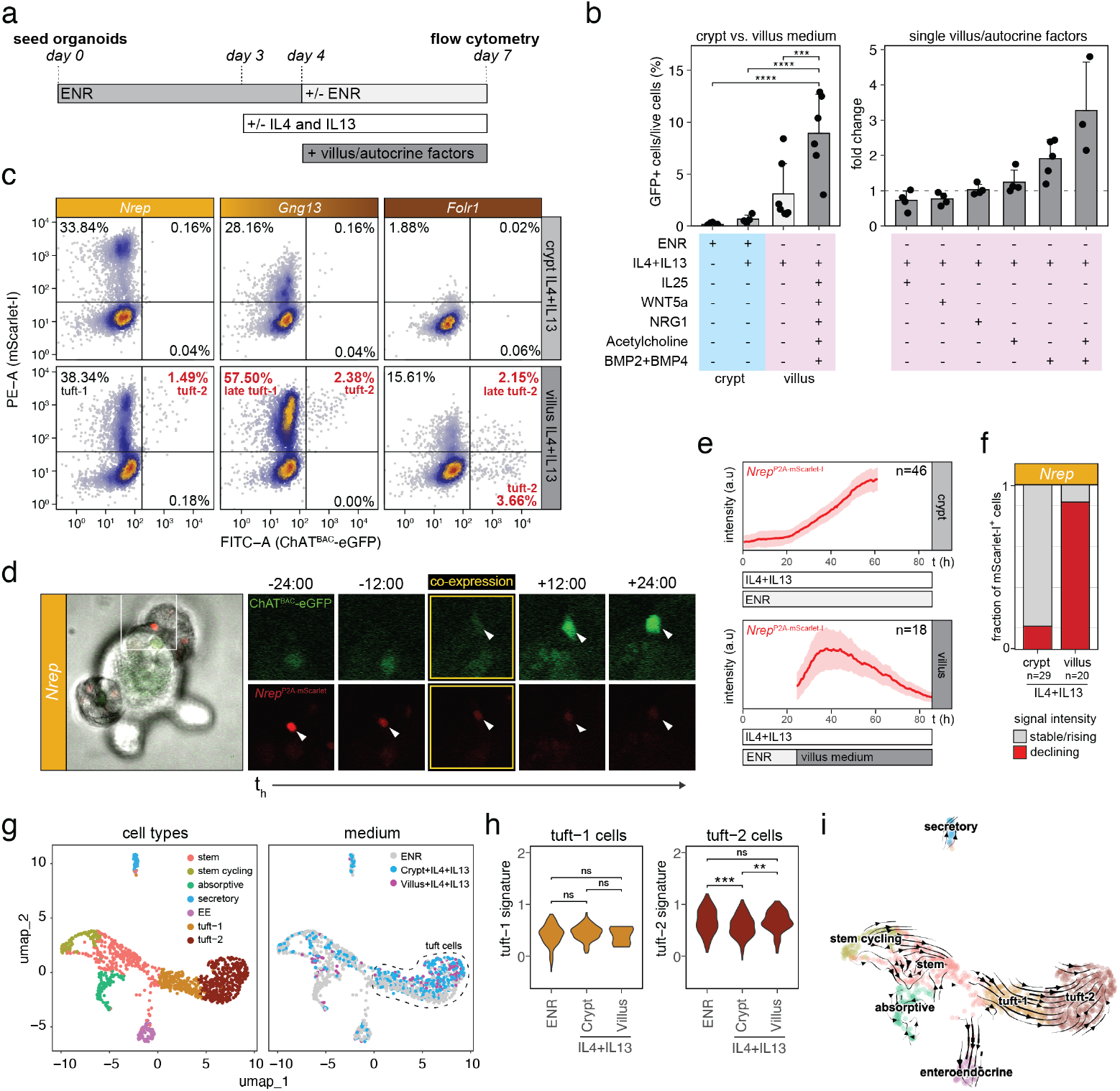
Crypt-villus signaling gradients facilitate transitioning from tuft-1 to tuft-2 states. **(a)** Culture setup to test tuft cell differentiation. Tuft-type reporter organoids were pretreated with IL-4 and IL13 on the 3^rd^ day after seeding in ENR (EGF, R-spondin and Noggin) medium. On the 4^th^ day, various factors were added to, or depleted from the medium. Frequency of tuft-type induction was tested at day 7 with flow cytometry. **(b)** Left, percentage of ChAT-eGFP^+^ cells in organoids treated with IL-4 and IL-13 in presence or absence of crypt or villus/autocrine-inspired signaling factors. Right, fold change induction of ChAT-eGFP^+^ cells after addition of single villus/autocrine-inspired factors to cytokine-pretreated organoids without ENR. Data are represented as mean ± SD (ANOVA p < 0.0001, *** adjusted p-value < 0.001, **** adjusted p-value < 0.0001, Tukey HSD test). **(c)** Representative flow cytometry analysis of fluorescent population frequencies in indicated tuft-type reporter organoids treated with ENR and cytokines (crypt+IL4+IL13) or villus-inspired medium (no ENR; with IL-25, WNT5a, NRG1, BMP2, BMP4 and acetylcholine chloride) after IL-4 and IL-13 pretreatment. **(d)** Microscopic stills from live imaging experiments of organoids in villus-inspired medium (no ENR; with IL-25, WNT5a, NRG1, BMP2, BMP4 and acetylcholine chloride) after IL-4 and IL-13 pretreatment. Left: confocal brightfield image of ChAT^BAC^-eGFP;*Nrep*^P2A-mScarlet-I^ organoid at the start of imaging. Right: stills of fluorescence channels (mSarlet-I: red, eGFP: green) from timepoints preceding and following co-expression of fluorescent markers. Cell that shows co-expression is indicated with a white arrowhead throughout imaging timepoints. **(e)** Average mScarlet-I signal over time in *Nrep*^P2A-mScarlet-I^ organoids in medium with ENR and cytokines (top; crypt) or villus-inspired medium (no ENR; with IL-25, WNT5a, NRG1, BMP2, BMP4 and acetylcholine chloride) after IL-4 and IL-13 pretreatment (bottom; villus). Shading in plot represents SEM. Number of cells comprising each graph are indicated. **(f)** Barplot showing fraction of mScarlet-I^+^ cells in *Nrep*^P2A-mScarlet-I^ organoids with stable/rising fluorescence signal or declining fluorescence signal in crypt or villus medium with cytokines (IL4+IL13). Related to e. Number of cells comprising each graph are indicated. **(g)** Transcriptomic analysis of single cells in organoids upon induction with cytokines in ENR or villus-inspired culture conditions. Left, UMAP of organoid cells being enriched for tuft cells by FACS. Cell types are annotated by color. EE: enteroendocrine cells. Right, same UMAP but cells are colored by medium condition. **(h)** Violin plots depicting distribution of relative levels of tuft signature scores in tuft-1 cells (left) or tuft-2 cells (right) origating from different culture conditions (ns not significant, ** p-value < 0.01, *** p-value < 0.001, Wilcoxon rank sum test). **(i)** RNA velocity-based trajectory inference superimposed on UMAP of Fig. 5i predicts unidirectional differentiation from tuft-1 to tuft-2 transcriptomic states.

To test the effect of our villus-inspired maturation medium on the transitioning of tuft-1 to tuft-2 phenotypes, we evaluated their frequencies in our tuft type-specific reporter organoids. This showed a clear shift in the maturation status of the tuft cell populations, with an increase in *Gng13*+ (late) tuft-1 cells, but in particular the emergence of mature tuft-2 cells expressing *ChAT* and *Gng13* or *Folr1* (Fig. 5c). Importantly, similar to cytokine-induced differentiation in ENR conditions, we used real-time imaging to confirm that the tuft-2 cells originated from Nrep^+^ tuft-1 cells in villus-inspired maturation medium (Fig. 5d). Moreover, under these villus-mimicking conditions, mScarlet-I signal in *Nrep*^P2A-mScarlet-I^ reporter organoids showed a declining profile after reaching peak intensity (Fig. 5ef), recapitulating *in vivo* expression patterns (Fig. 3f). This contrasts with standard culture conditions reminiscent of a crypt environment, where we mostly observed stabilizing mScarlet-I signals at their peak intensity (Fig. 5ef).

Lastly, we performed scRNA-seq on reporter organoids treated with our villus-inspired maturation medium to confirm a shift in maturation states and no induction of alternative non-physiological transcriptomic states. Unsupervised clustering with our previous organoid dataset showed mixing of tuft cells derived from the different medium conditions on the UMAP (Fig. 5g). Furthermore, tuft-1 and tuft-2 signature scores were similar in the crypt and villus medium conditions (Fig. 5h). This indicates that, while the maturation medium had strong effects on the frequency of tuft-2 cells (Fig. 5bc), it did not induce tuft cell transcriptomic states that were previously not observed. Moreover, trajectory inference through RNA velocity analysis corroborated the unidirectional differentiation trajectory from tuft-1 to tuft-2 transcriptomic states, which we repeatedly observed in our live imaging experiments (Fig. 5i). Collectively, these experiments show that crypt-villus signaling gradients control the transitioning of tuft-1 to tuft-2 phenotypes, and its manipulation can be used to increase the frequency of mature tuft-2 phenotypes in organoids.

### An organoid-based platform for functional characterization of tuft cell properties

Our tuft-type reporter organoids in combination with the optimized tuft maturation medium provide a powerful *in vitro* platform to study the basic cell biology that governs mature tuft cell functioning. To demonstrate its utility, we first set out to investigate the dynamics of the typical bottle-shaped tuft cell morphology^26^ (Fig. 6a), which features distinctive membrane characteristics. In addition to the well-known robust protrusions at the apical surface that form the tuft of the tuft cell, EM studies have also identified the presence of lateral microvilli as well as large basolateral extensions^27^, which are speculated to be involved in cell-cell communication with neighboring cells^26,28–32^. Within our live-cell imaging recordings, we indeed observed long protrusions at the basolateral membrane of tuft cells (Fig. 6b). Unexpectedly, these protrusions were highly dynamic, revealing growth and shrink rates of more than 10 μm per hour (Fig. 6c). Moreover, long protrusions (∼20μm, up to 6 hours lifetime) showed more stability than short ones (5μm, one hour lifetime) (Fig. 6d). Next, imaging Chat^+^ tuft-2 cells in combination with a nuclear marker confirmed that these dynamic protrusions were indeed basolateral extensions (Extended Data Fig. 6a). Moreover, also Nrep^+^ tuft-1 cells are capable of generating such basal protrusions (Fig. 6e). These novel experimental opportunities highlight how future studies may improve our limited understanding of tuft cell morphological changes and its relation to functioning.

**Figure 6:**
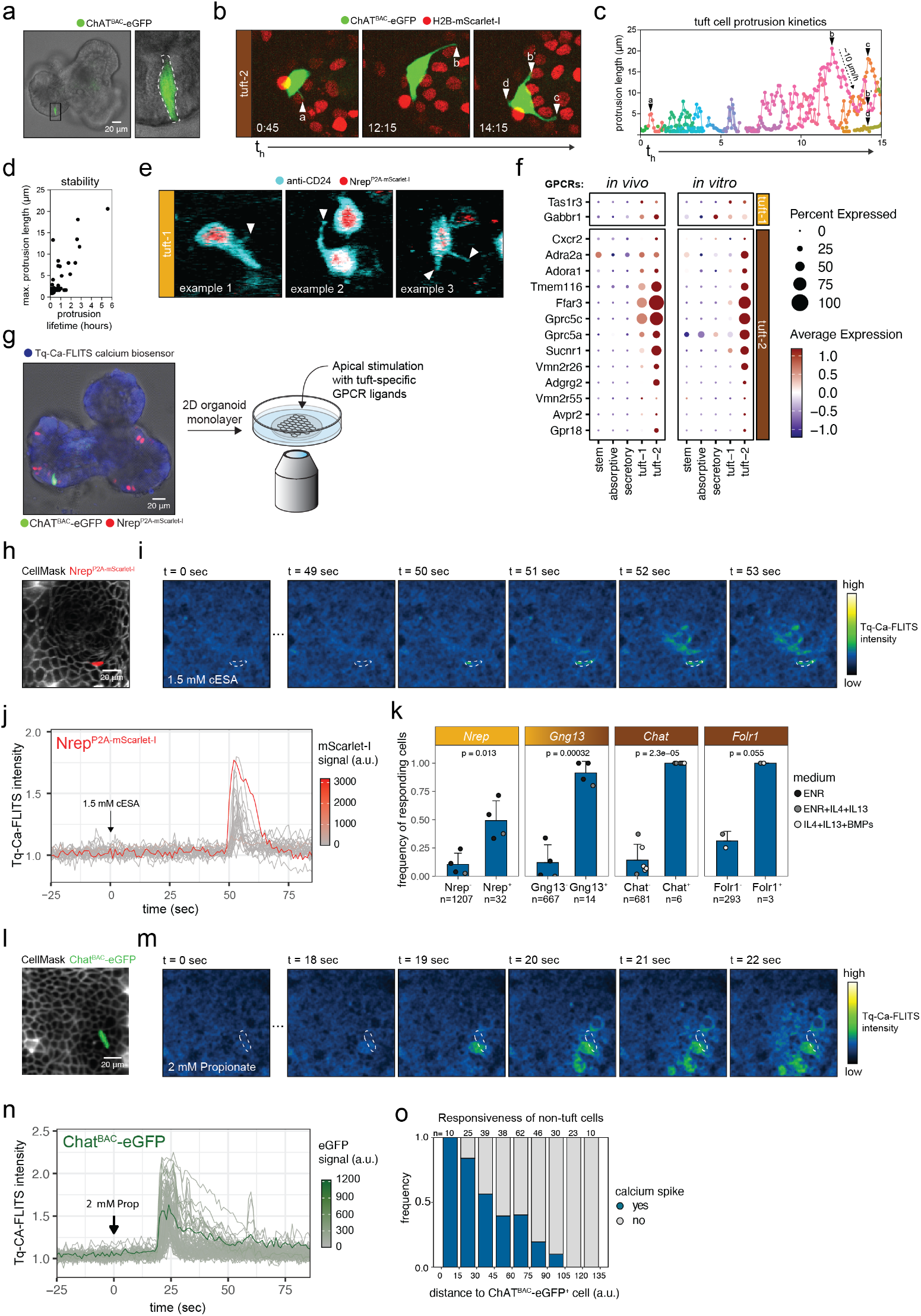
An organoid-based platform for functional characterization of tuft cell properties. **(a)** Microscopic images of ChAT^BAC^-eGFP cell in organoid showing typical tuft cell morphology. Left: overview confocal brightfield image overlayed with GFP fluorescence channel of whole organoid. Right: zoom-in of indicated region. White dotted line outlines ChAT^BAC^-eGFP tuft cell. **(b)** Live imaging of ChAT^BAC^-eGFP cells in organoids reveal dynamic protrusions in ChAT^+^ tuft-2 cells that can span several neighbouring cells. Merge of fluorescence channels is shown: green, eGFP; red, nuclei visualized with H2B-mScarlet-I. Three stills of indicated timepoints are shown. White triangles with letters indicate protrusions. **(c)** Protrusion lengths change rapidly over time (in the order of 10 μm/h). Triangles with letters indicate single protrusion tracks corresponding to protrusions shown in microscopy images of a. **(d)** Relationship between maximum protrusion length and protrusion lifetime shows longer protrusions to be more stable. **(e)** Microscopic stills from live imaging of *Nrep*^P2A-mScarlet-I-nls^ organoids stained with anti-CD24, a general and early marker for secretory cells, to visualize cell membranes and protrusions of Nrep^+^ tuft-1 cells. Merge of fluorescence channels is shown: cyan, anti-CD24; red, *Nrep*^P2A-mScarlet-I-nls^. White triangles with letters indicate protrusions. Stills of three different examples of Nrep^+^ cells with dynamic protrusions are shown. **(f)** Relative expression level of tuft-specific GPCRs within indicated cell types, as extracted from the mouse *in vivo* integrated single-cell dataset (fig. 1g) and organoids (fig. 3h). Average expression is represented by the dot color while the percentage of expressing cells is denoted by the dot size. **(g)** Experimental setup for live-cell imaging experiments to monitor tuft cell activation. Tuft-type reporter organoids expressing the Tq-Ca-FLITS calcium biosensor are re-plated to form 2D monolayers that can be apically stimulated and are compatible with high temporal resolution imaging. **(h)** Fluorescent image of 2D monolayer of reporter organoid with CellMask deep red incubation to label cell boundaries (membranes, in white) and mScarlet-I reporter signal to label tuft identity. Current image is prior to stimulation with cis-epoxysuccinic acid (cESA). **(i)** Fluorescent intensity fluctuations of Tq-Ca-FLITS biosensor within the field-of-view cell monolayer as shown in panel d, following stimulation with cESA at t=0. Time (seconds) post cESA exposure is indicated at the top of each panel. White outline indicates Nrep^+^ tuft cell. **(j)** Single-cell traces of Tq-Ca-FLITS intensity fluctuations within monolayer of panel e. Traces are colored by mScarlet-I signal measured in the corresponding cell. **(k)** Responsiveness of tuft-reporter positive cells following stimulation with 1.5 mM cESA. Per bar, each point represents a separate experiment and is colored for the medium used. Data are represented as mean ± SD (number of cells comprising each bar and p-values are indicated, t-test). **(l)** As in panel d, but now monolayer of ChAT^BAC^-eGFP reporter organoid prior exposure to propionate. **(m)** As in panel e, but Tq-Ca-FLITS intensity fluctuation post propionate exposure. Time since propionate addition is indicated at the top of each panel. Arrow indicates ChAT^BAC^-eGFP^+^ tuft cell. **(n)** Single-cell traces of Tq-Ca-FLITS intensity fluctuations within monolayer of panel i. Traces are colored by GFP signal measured in the corresponding cell. **(o)** Responsiveness of non-tuft cells (mScarlet-I^-^ GFP^-^) in the proximity of the ChAT^BAC^-eGFP^+^ cell shown in h. Number of cells comprising each bar are indicated at the top of each bar. Results of one representative experiment are shown.

Next, we investigated tuft cell chemosensing that, similar to cells in the taste buds, involves GPCR-mediated signaling to control intracellular calcium pulses and subsequent membrane depolarization. Therefore, we complemented our tuft-type reporter organoids with a genetically-encoded calcium biosensor (Tq-Ca-FLITS^33^), to visualize and measure the GPCR-mediated activation potential of putative stimuli (Extended Data Fig. 6b). We extracted tuft-specific GPCRs from our high-resolution integrated *in vivo* scRNA-seq dataset (Fig. 6f). Reassuringly, the resulting list contained GPCRs that were previously found to be tuft-specific, such as *Sucnr1* and *Vmn2r26*, as well as some novel tuft-specific GPCRs, like *Gabbr1* and *Gpr18*. Most of the tuft-specific GPCRs recapitulated their *in vivo* expression pattern in organoids, with expression generally being highest in tuft-2 cells (Fig. 6f).

To develop a screening setup that enables apical stimulation and single-cell measurements of calcium influx in real-time, we converted calcium biosensor-expressing reporter organoids into 2D monolayers^34,35^ [see Methods section] (Fig. 6g). We first exposed these monolayers to the succinate mimetic cis-epoxysuccinic acid (cESA), a known stimulus (Fig. 6h). As expected, we observed calcium spikes within a minute after administration of the ligand (Fig. 6i). While tuft cells were the first to respond to the stimulus (Fig. 6j), calcium spikes did not remain exclusive to tuft cells and were similar in strength between tuft and non-tuft cells (Extended Data Fig. 6c). This propagation of calcium pulses to nearby non-tuft cells is analogous to what has recently been observed in the trachea^36^. Concordant with the highest *Sucnr1* expression in tuft-2 cells (Fig. 6f), we observed the highest response probability in cells positive for tuft-2 markers (Fig. 6k). As a proof-of-principle of our functional platform to study chemosensing and screen for novel stimuli, we next tested responsiveness to the *FFAR3* ligand propionate, a microbiota-derived short chain fatty acid which is speculated but never formally shown to act directly on tuft cells. Stimulation with propionate readily triggered calcium spikes in mature ChAT^+^ tuft cells (Fig. 6l and Fig. 6m), which propagated to nearby non-tuft cells (Fig. 6n and Fig. 6o). Unlike stimulation with the succinate-mimetic cESA, Nrep^+^ tuft-1 cells showed no response to propionate (Extended Data Fig. 6d), underscoring the importance of studying chemosensing in fully mature functional tuft-2 cells.

## Discussion

Tuft cells in the intestine function as epithelial orchestrators of host defense against luminal parasites^5–7,17^. Beyond their role during infections, alterations in intestinal tuft cell numbers and function are associated with immune-related diseases such as inflammatory bowel disease, coeliac disease and neoplasias^37^. Despite scientific and clinical interest in tuft cells, their rare occurrence *in vivo* and the absence of accurate *in vitro* models, have proven to be practical limitations that hamper investigation of the cell biological processes underlying tuft cell functioning. Here, we unlock organoids as an experimental model system to study tuft cell functioning *in vitro*. We show that the various transcriptomic states of tuft cells *in vivo* and their interdependencies are truthfully reflected in organoids, optimize culture conditions to enrich for the most mature tuft cell states, and showcase experimental opportunities to study the dynamics of distinctive tuft cell morphology, behavior and chemosensing properties at high resolution.

In our in-depth characterization of *in vivo* tuft cell phenotypes, upfront enrichment of ChAT^+^ cells prior to single-cell transcriptomic profiling ensured a substantial number of high-quality tuft cell profiles. In addition to confirmation of the earlier reported tuft-1 and tuft-2 transcriptomic states within the tuft cell pool^14^, our data revealed frequent presence of intermediate tuft cell profiles with co-expression of both tuft-1 and tuft-2 markers. This continuum of transcriptomic states, with no clear demarcation between the two previously described tuft-1 and tuft-2 types, argues against an early binary confinement to either of two mature tuft cell types during differentiation. Using tuft type-specific reporter knock-ins in organoids, we investigated directionality over the spectrum of tuft profiles by real-time tracking of cell identity and showed that cytokine-induced tuft cell differentiation follows a linear trajectory from a tuft-1 to a tuft-2 state. This indicates that the tuft-1 and tuft-2 transcriptional programs are expressed sequentially within the same cell and not by two separate mature tuft types and explains the earlier reported spatial preference of the tuft-1 and tuft-2 cells to the crypt and villus compartments, respectively^15^. Previous timelapse imaging of organoids from a dual *Trpm5*^GFP^:*Adgrg2*^mCherry^ reporter mouse, in combination with histological analysis of intestinal tissues, also showed emerging expression of tuft-2 marker *Adgrg2* in cells that already expressed general tuft cell marker *Trpm5*, providing independent support for our observations^38^.

Using our organoid tuft reporters, we show that tuft cell maturation is stimulated by BMP signaling, which also supports functional zonation of enterocytes, goblet cells and enteroendocrine cells along the crypt-villus axis^25,39^. Previously, BMP2 signaling originating from tuft cells was shown to prevent *de novo* tuft cell specification from stem cells located at crypt bottoms^23^. This does not contradict our finding that BMP ligands, increasingly expressed by the intestinal stroma towards villus tips, promote maturation of already existing tuft cells. Similarly, microbiota-derived butyrate has been shown to limit tuft cell specification from stem cells^22^, but its effect on tuft cells that populate villi is unknown.

While ChAT reporter mice have been confirmed to label all cells committed to the tuft lineage ^17^, our transcriptomic datasets and organoid timelapse measurements indicate that ChAT expression levels increases during tuft cell maturation. Even though we included cells with minor ChAT expression levels in our analyses, our study mainly provides insights into the final steps of tuft cell differentiation and maturation. In contrast, many studies have focused on the initial steps of stem cell specification towards the tuft lineage^9–13^, and recently the capacity of tuft cells to act as reserve stem cells^40^. While our data is less suited to investigate early fate decisions, we did observe a significant fraction of tuft cells with a progenitor phenotype. These tuft-p cells expressed markers of proliferative states as well as markers of other epithelial lineages, hinting at recent multilineage potential. Consequently, single marker genes specific to the tuft-p cells were not identified. This prohibited their genetic labeling with knock-in reporters and has led to an incomplete understanding of how tuft-p cells fuel into the more advanced, post-mitotic stages of tuft cell differentiation. We found few phenotypic traits in tuft-1 cells that are unique to the tuft lineage, yet absent in tuft-2 cells. Rather they seem to express most tuft-specific genes at a lower level when compared to tuft-2 cells, akin to a more immature state. Of the few tuft-specific genes that showed higher expression in tuft-1 than in tuft-2, *Nrep* was the most prominent marker being exclusive to tuft-1 cells in both mice and human. Alternatively, most tuft-specific genes, and in particular GPCRs that play a central role in chemosensing as well as immune effector molecules, showed increasing expression levels from tuft-1 to tuft-2 in both mouse and human. Furthermore, our organoid models allowed us to show that both tuft-1 and tuft-2 cells generate highly dynamic basolateral protrusions of several tens of microns long. For future studies, it will be intriguing to investigate how diverse input signals are processed by tuft cells, how they are communicated to other cells, and the relationship between dynamic morphology and function.

Understanding the unique cell biology that underlies tuft cell functioning is of high interest to identify therapeutic targets to intervene with immune-related disorders and infections in the intestinal tract. The findings of our study show high similarity between the mature tuft cell phenotypes in mice, organoids and human. Moreover, mouse organoids are fully complementary with *in vivo* models and will be instrumental to generate mechanistic insights that can then be tested in physiological settings. Our comprehensive characterization and comparison of tuft cell phenotypes *in vivo* and in organoids, paired with reporter models and optimized differentiation strategies, established a functional platform to address the poorly understood mechanisms of perception, processing and transmission of chemosensory signals by tuft cells.

## Acknowledgements

We gratefully acknowledge all lab members for reagents, suggestions, and discussions and the UMCU Flow Core Cytometry Facility for support. We thank Prof. Dr. Gadella (van Leeuwenhoek Centre for Advanced Microscopy, Swammerdam Institute for Life Sciences, University of Amsterdam) for providing the Tq-Ca-FLITS plasmid. This work is funded by the ‘Organoids in time’ (OCENW.GROOT.2019.085) and Gravitation programme (IMAGINE!; 024.005.009) from the Netherlands Organization for Scientific Research (NWO). This work is part of the Oncode Institute, which is partly financed by the Dutch Cancer Society. This paper was typeset with the bioRxiv word template by @Chrelli: www.github.com/chrelli/bioRxiv-word-template

## Author contributions

Conceptualization, J.R.B.d.A. and H.J.G.S.; Methodology, J.R.B.d.A., J.H. and R.H.; Investigation, J.R.B.d.A., J.B., M.A.B., T.V., I.V-K., M.H., I.J. and

D.L.; Formal analysis, J.R.B.d.A. and M.A.B.; Data curation, S.R.B.; Software, M.A.B.; Visualization, J.R.B.d.A. and M.A.B.; Writing – original draft,

J.R.B.d.A. and H.J.G.S.; Resources, M.G.; Supervision, H.J.G.S., J.S.Z. and S.J.T.; Funding acquisiton, H.J.G., J.S.v.Z., S.J.T., J.v.R. and H.C.

## Materials and Methods

### Mice

*ChAT*^BAC^-eGFP mice were obtained from the Jackson Laboratory and housed at the Hubrecht institute and Netherlands Cancer Institute mouse facilities under specific pathogen-free (SPF) conditions. For all experiments, adult mice at least 6 weeks of age, of either sex, were used.

### Organoid derivation and culture

Small intestinal crypts were isolated as described previously^41^. In brief, mouse small intestines were excised, rinsed with PBS and incubated with DPBS containing 5 mM EDTA for 40 min at 4 °C on a carousel to extract crypts. Crypts were pelleted by centrifugation of the supernatant, washed with advanced DMEM (Gibco) and plated in drops of Matrigel (Corning) to form organoids. Organoids were derived and cultured with ENR medium: advanced DMEM/F12 (Gibco) supplemented with 1% Penicillin/ Streptomycin (P/S, Lonza), 1% HEPES buffer (Gibco) and 1% GlutaMAX (Gibco), 5% R-spondin conditioned medium (in-house production), 10% Noggin conditioned medium (in-house production), B27 (Invitrogen), 1.25 mM N-acetylcysteine (Sigma-Aldrich) and 50 ng/ml EGF (Invitrogen). Organoids were maintained at 37 °C with 5% CO_2_ and passaged weekly through fragmentation by shear-stress (pipetting). Medium was refreshed every 2 days.

### Organoid tuft cell differentation

3D organoid cultures were subjected to differentiation regimens as indicated. In general, tuft cell specification was induced on the 3^rd^ day after passaging with ENR medium supplemented with recombinant murine IL-4 and IL-13 (10 ng/ml each, Immunotools). Thereafter, tuft cell maturation was facilitated by removal of EGF, Noggin and R-spondin from the medium and addition of one of the following or a combination thereof on the 4^th^ day after passaging: recombinant murine IL-25 (20 ng/ml, Immunotools), recombinant human BMP2 (20 ng/ml, Immunotools), recombinant human BMP4 (20 ng/ml, Immunotools), recombinant human NRG1 (20 ng/ml, R&D systems), recombinant murine WNT5a (20 ng/ml, Biotechne) and acetylcholine chloride (100 μM, Sigma Aldrich). Organoids were harvested for downstream analysis on the 7^th^ day after passaging.

### Tissue and organoid dissocation for flow cytometry and sorting

For isolation of tuft cells from mouse tissue, small intestines were excised, rinsed with PBS and incubated for 10 min in PBS containing 100 μM DTT. Next, the small intestines were cut into smaller fragments, transferred into DPBS containing 5mM EDTA and spun on a carousel for 40 min at 4 °C. Organoids were extracted from Matrigel with cold advanced DMEM. To make single-cell suspensions, tissue fragments or organoids were trypsinized for 5 min at 37 °C with TypLE containing 10 μM Y-27632 Rho-kinase inhibitor to inhibit anoikis. Trypsinized single-cell preparations were washed with advanced DMEM, filtered (40 μm cell strainer) and resuspended in advanced DMEM containing 10 μM Y-27632. In case of immunofluorescence, single-cell suspensions were stained for 30 min on ice with EpCAM APC (Clevers lab) 1:100 and CD45 AF700 1:100. Cells were stained briefly with DAPI prior to flow cytometry.

### Flow cytometry

Single-cell suspensions were prepared as described above. Flow cytometry measurements were performed on a BD FACSCelesta CellAnalyzer. Single live cells (DAPI^-^) were gated and GFP and mScarlet-I fluorescence was measured in the FITC-A and PE-A channels, respectively. Gates were set based on negative control samples. Flow cytometry data was analyzed and visualized using BD FACSdiva software and the flowCore (2.14.0) and CytoExploreR (1.1.0) R packages.

### Plate-based scRNA-seq

Single-cell suspensions were prepared as described above. Viable single cells (DAPI^-^) were sorted (BD FACSAria III) into 384-well cell-capture plates from Single Cell Discoveries, which contain a 50 nl droplet of well-specific barcoded primers and 10 μl of mineral oil (Sigma M8410). After sorting, plates were briefly centrifuged at 500 × *g* and then kept on dry ice till further storage at –80 °C. Single-cell RNA sequencing was performed by Single Cell Discoveries according to an adapted version of the SORT-seq protocol^42^ with primers described in van den Brink *et al*. ^43^ Cells were heat-lysed at 65 °C followed by cDNA synthesis. After second-strand cDNA synthesis, all the barcoded material from one plate was pooled into one library and amplified using in vitro transcription (IVT). Following amplification, library preparation was performed following the CEL-Seq2 protocol^44^ to prepare a cDNA library for sequencing using TruSeq small RNA primers (Illumina). The DNA library was sequenced by paired-end sequencing on an Illumina Nextseq™ 500, high output, with a 1×75 bp Illumina kit (read 1: 26 cycles, index read: 6 cycles, read 2: 60 cycles).

### scRNA-seq analysis

For alignment of reads, an adopted version of the nf-core scrnaseq pipeline (2.4.0)^45^ was used (https://github.com/gowanaka/nf-core-scrnaseq). In brief, STARsolo (2.7.10b) was used to align reads to a custom mm10 transcriptome including *eGFP* and *P2A-mScarlet-I-P2A-DTR* transgenes and ERCC spike-ins. Following mapping, count matrices were generated with STARsolo (2.7.10b). Gene expression was analysed using Seurat (5.0.1)^46^. Cells with <30% mitochondrial content, <25% exogenous ERCC spike-in content and >1000 detected genes were selected for downstream analysis. Mitochondrial transcript counts were removed prior to count normalization and scaling by the Seurat NormalizeData and ScaleData functions, respectively. Seurat CCA-integration was performed to eliminate technical plate-based batch effects and subsequently unsupervised clustering was used to cluster cells according to the standard Seurat workflow. Gene expression signature scores were calculated with the Seurat AddModuleScore function. Differential expression analysis was performed with the FindAllMarkers function. Tuft specificity scores for Figure 3c were calculated as log_10_((1/average expression in non-tuft epithelial cells)+1). Integration with published single-cell transcriptomic datasets (Haber *et al*. GSE92332, Zwick *et al*. GSE201859 and Böttcher *et al*. GSE152325) was performed with the Seurat package (harmony integration) according to the standard workflow. Clusters were annotated according to the cell type annotations included with the published datasets. GO-term enrichment analysis was performed with clusterProfiler (4.8.3)^47^. RNA velocity analysis was performed with scVelo (0.3.2)^48^.

### Analysis of published RNA sequencing data

Transcript counts from mice infected with *Nippostrongylus brasiliensis* and *Bmpr1a* knockout mice were obtained from E-MTAB-9183^23^ and GSE194004^25^, respectively. The R package DESeq2 (1.40.2)^49^ was used to perform differential expression analysis, followed by gene set enrichment analysis (GSEA) with clusterProfiler (4.8.3)^47^ and visualization with enrichplot (1.20.3).

### Generation of organoid knock-ins

Organoid knock-ins were generated by in-trans paired nicking (ITPN) as described in Bollen and Hageman *et al*^24^ (Extended Data figure S3cd). Cas9 D10A nickase (addgene #48141) locus-specific expression vectors were generated according to published protocols^50^. For transfection, organoids were trypsinized to cell clumps containing ∼5 cells (1×10^6^ cells total) and coelectroporated with 15 μg DNA and 5 ug of Cas9 D10A nickase and targeting vector using the NEPA21 Super Electroporator (Nepagene) following described conditions^51^. Electroporated cell clumps were plated in Matrigel overlayed with ENR culture medium, which was supplemented with 10 μM Y-27632 Rho-kinase inhibitor and 0.25 nM Wnt surrogate-FC fusion protein (U-Protein Express) for the first 3 days. Targeted cells were selected using 2 μg/ml puromycin and maintained as polyclonal populations. Proper integration of the knock-in constructs was validated by targeted PCR of genomic organoid DNA (isolated with QIAamp DNA Micro Kit from Qiagen, according to the manufacturer’s instructions), followed by Sanger sequencing of the PCR product using the following primers:

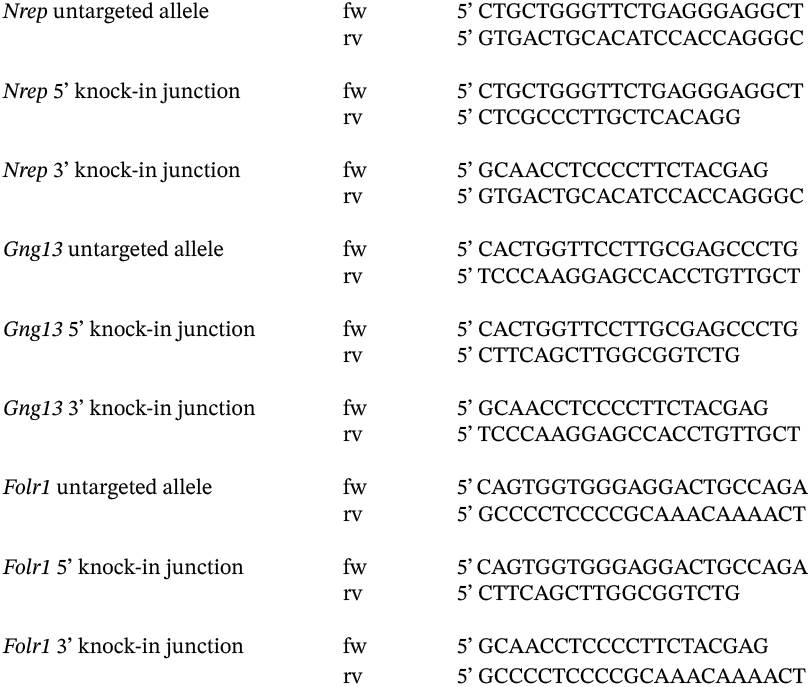

### Lentiviral transduction of organoids

Organoids were transduced with lentivirus encoding hEF1a-Tq-Ca-FLITS or H2B-mScarlet-I followed by an IRES and a blasticidin resistance cassette. In brief, organoids were incubated in trypsin at 37 °C to make a suspension of cell clumps containing approximately 5-10 cells. Cell clumps were transduced by spinoculation for 1h at room temperature and then plated in Matrigel overlayed with ENR culture medium, which was supplemented with 10 μM Y-27632 Rho-kinase inhibitor and 0.25 nM Wnt surrogate-FC fusion protein (U-Protein Express) for the first 3 days. Transduced organoids were selected using 3 μg/ml blasticidin (InvivoGen) and maintained as polyclonal populations.

### Calcium imaging of 2D organoid monolayers

For calcium imaging of 2D monolayers, organoids were fragmented and seeded on polyacrylamide gels (18 kPa stiffness) that had been coated overnight with 100 μg/ml Laminin [Sigma-Aldrich L2020] and 250 μg/ml Collagen I [First Link (UK)] as in Pérez-González *et al*^34^. After seeding, organoid fragments were overlayed with ENR medium (described above), which was supplemented with 1x N2 Supplement [Thermo Scientific], 10 ng/ ml human-FGF2 [Peprotech], 3 μM Chiron [Bio-Connect], 10 μM Y-27632 and 100 ug/ml primocin [InvivoGen], and contained a higher amount of R-spondin conditioned medium (20%). Chiron and Y-27632 were removed from the medium one day later. For differentiation of 2D monolayers, IL-4 and IL-13 (10 ng/ml each) were added 2 days before imaging, followed by addition of BMP2 and BMP4 (20 ng/ml each) in combination with removal of Noggin from the medium on the day prior to imaging. Timelapse imaging was performed 3-4 days after seeding of organoid crypts, when monolayers formed mature crypt-villus compartments, on a Leica SP8 WLL scanning confocal microscope (40x-water-N.A.1.1, 37 °C, 6% CO_2_) with Leica Application Suite X at 1 frame per second using resonant scanning (8000 Hz). The genetically encoded Tq-Ca-FLITS^33^ calcium biosensor was excited with an ultraviolet laser at 405 nm, to minimize bleed through of EGFP in the Tq-Ca-FLITS channel, and emission was measured at 445-505 nm. Organoid monolayers were stimulated during imaging with cis-epoxysuccinic acid (cESA, 1.5 mM, Fischer Scientific) or propionate (2 mM, Sigma-Aldrich P1880), as indicated. CellMask Deep Red (1:20000, Invitrogen, 640 nm excitation and 655-710 nm emission) was added 30 min prior to imaging to stain plasma membranes and enable post-acquisition cell segmentation. Before acquisition of each timelapse, EGFP (488 nm excitation and 495-545 nm emission) and mScarlet-I (570 nm excitation and 590-630 nm emission) signals were measured to determine tuft reporter status per cell in the field of view.

### Live organoid timelapse imaging

For live imaging, organoids were mechanically fragmented and seeded in cold BME (Trevigen) in a four-well chambered cover glass (#1.5 high-performance cover glass, Cellvis). After seeding, wells were placed on a cold block for 10 minutes allowing the organoids to sink to a position close to the glass, as described earlier^52^. Organoids were cultured for 2 to 3 days in ENR medium until clear crypt-villus structures were apparent. Next, IL4 and IL13 (10 ng/ml each) were added to the medium and organoids were imaged for 3 days, starting on the 1^st^ or 5^th^ day after interleukin treatment to capture transitioning fluorescence phenotypes of all tuft reporters. Imaging was performed at 37 °C and 5% CO_2_ with a scanning confocal microscope, using either the Leica TCS SP8 with a 40× water immersion objective (numerical aperture, 1.10) or the Nikon A1R MP with a 40x oil immersion objective (NA = 1.30). The voxel size used was 0.4×0.4×2 micron for both microscopes. The maximum time resolution used was 24 minutes per frame.

### Timelapse analysis

All tracking and timelapse analyses were performed using OrganoidTracker 2.0^53^. To measure marker intensities the average fluorescent intensity around the nucleus center was measured using a Gaussian kernel with a width of 1.5 pixels. To avoid measuring bleed through from the green (ChAT^BAC^-eGFP) channel we subtracted the green channel from the red channel (all other markers) such that there was no signal in the cytoplasm, where Nrep/Gng13/ Folr1^P2A-mScarlet-I-nls^ marker signal should be absent. Tuft cell locations were determined by evaluation of transmission signal. Cells in the spherical end of organoid buds were categorized as located in the crypt, while cells that fitted into a sphere drawn around the central villus region were deemed to be in the villus. Cells in the narrow region between crypt and villus compartment were deemed to be in the neck region. The crypt and the neck region were combined when analyzing the colocalization of ChAT and DCLK1 (Fig. 3d). For the analysis of the relative timing of marker expression and the locations of emergence, data from at least three replicates was pooled. For the *Nrep*^P2A-mScarlet-I^ signal in villus-inspired and standard crypt medium, a representative organoid was analyzed.

### Automated tuft cell protrusion analysis

To track protrusions, we first extracted a 3D cell mask by thresholding on the ChAT^BAC^-eGFP signal. We then performed binary erosion and dilation operations to extract the mask of the cell body. The difference between the cell body and the full cell mask gave us the protrusions. Protrusion tips were defined by the pixel in the protrusion that was furthest away from the cell body. Protrusions were linked by connecting protrusions to the protrusion with which they shared maximum overlap in the next frame. If there was little or zero overlap (for instance if protrusions were very thin or small), protrusions were connected with the tips closest to each other. Using the nuclear marker, tuft cells were automatically rotated such that the epithelial plane was in XY. For this, we locally fitted an ellipsoid to the nuclear signal, so that the two major axes defined a plane tangential to the epithelial layer. These could then be used to find the proper rotation needed.

### Immunostaining of organoids

Immunofluorescence of organoids was performed as in Dekkers *et al*^54^. Briefly, organoids were harvested from Matrigel droplets with cold advanced DMEM/F12 (Gibco) supplemented with 1% Penicillin/Streptomycin (P/S, Lonza), 1% HEPES buffer (Gibco) and 1% GlutaMAX (Gibco) and pelleted by centrifugation. Following fixation with 4% paraformaldehyde in PBS at 4 °C for 45 min, organoids were permeabilized with 0.1% (vol/vol) PBS-Tween and blocked with PBS containing 0.2% bovine serum albumin and 0.1% Triton X-100 (4 °C for 15 min). Primary antibodies (anti-mouse CD45, 1:200, Biolegend 103102; anti-mouse Folr1, 1:100, R&D Systems AF6936-SP) were incubated overnight in blocking buffer at 4 °C with mild rocking. After washing in blocking buffer (4x), secondary antibodies (AF568 donkey anti-Sheep, Invitrogen A21099, 1:500; AF568 goat anti-rat, Invitrogen A11077, 1:500) were incubated, together with Hoechst 33342 (ThermoFischer Scientific 62249, 1:2000), in the same manner. Finally, organoids were washed in blocking buffer (4x) and mounted on glass slides in a fructose-glycerol clearing solution (60% (vol/vol) glycerol and 2.5 M fructose). Image acquisition was performed on a Leica SP8 WLL scanning confocal microscope (40x-water-N.A.1.1) with Leica Application Suite X.

### Immunostaining of tissues

Mouse small intestinal tissue stainings were perfomed as in *Snippert et al*.^55^ In short, intestines were fixed in 4% paraformaldehyde at room temperature for 20 min and washed in cold PBS. 1 cm^2^ of intestinal wall was put in a mold. Four percent low melting point agarose (40°C) was added and allowed to cool on ice. Once solid, a vibrating microtome was used to make semi-thick sections (150 μm). Sections were stained with anti-mouse b-catenin (Sigma C2206, 1:200), overnight at 4 °C with mild rocking. After washing, sections were incubated with AF568 goat anti-rabbit secondary antibody (Invitrogen A11036, 1:500) and counterstained with Hoechst 33342 (ThermoFischer Scientific 62249, 1:2000) Sections were embedded in Vectashield (Vector Laboratories) after a final washing step. Image acquisition was performed on a Leica SP8 WLL scanning confocal microscope (40x-water-N.A.1.1) with Leica Application Suite X.

### Quantification and statistics

Statistical analysis was performed as noted in figure legends using R (R base, ggplot2 (3.5.1), ggpubr (0.6.0) and Seurat packages (5.0.1)). All experiments were performed in multiple distinct replicates. Data distribution was assumed to be normal, but this was not formally tested. Statistical tests were two-tailed Student’s t-tests. For comparisons between more than two sample groups, one-way ANOVA was performed, using Tukey HSD for post-hoc analysis. Data are presented as mean +/-SD, unless otherwise stated in the figure legend.

### Data availability

Organoid and primary tissue single-cell RNA sequencing data presented in this study will be made publicly available upon publication.

## Supplementary information

**Supplementary table S1**

Differential expression analysis between three clusters of *ex vivo* ChAT^BAC^-eGFP scRNA-seq dataset (Wilcoxon Rank Sum test, p_adj_-value < 0.01). Relatad to Figure 1.

**Extended Data Figure 1.**
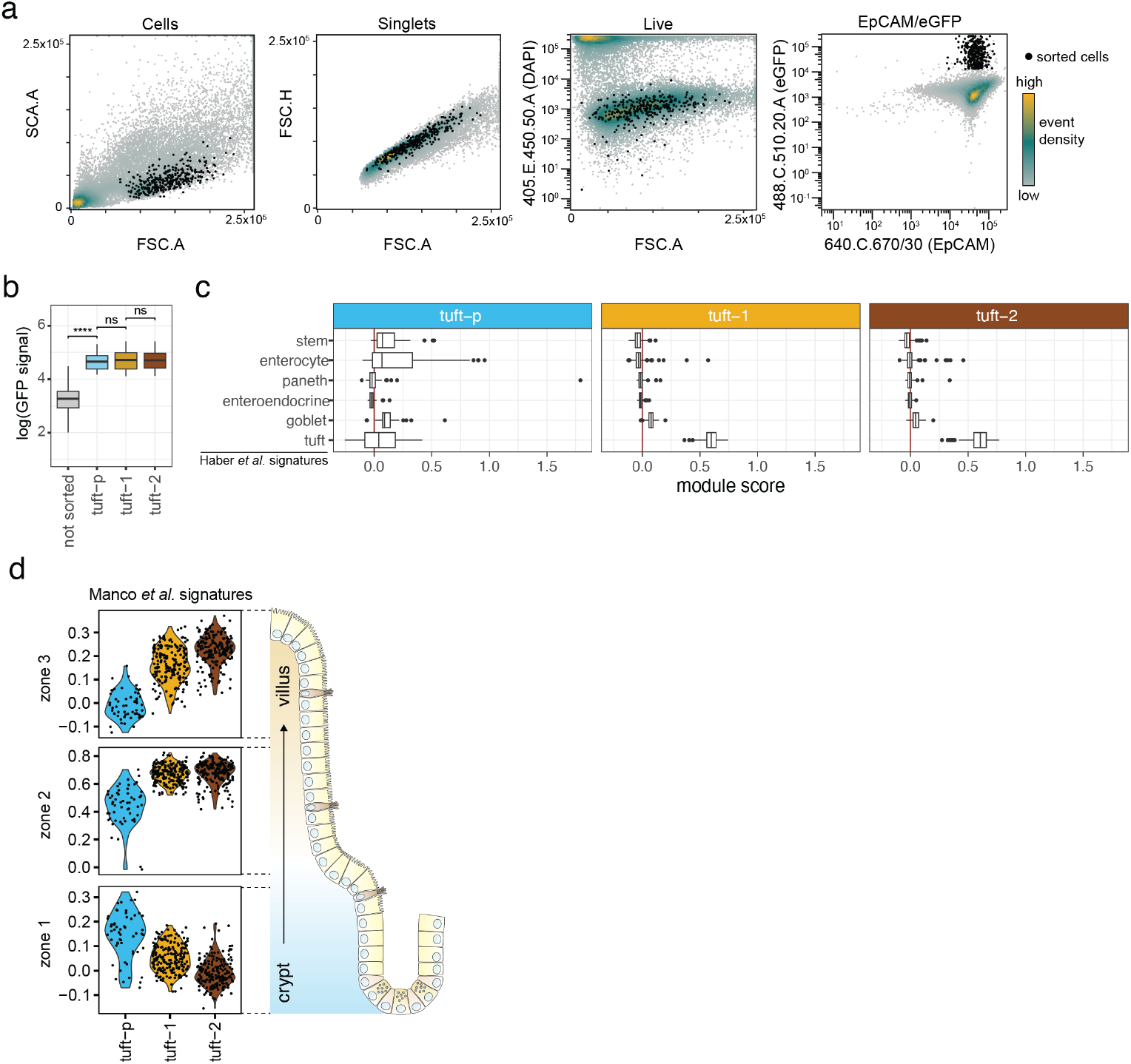
Small intestinal tuft cell heterogeneity and zonation Related to Figure 1. a. Gating strategy used during FACS prior to plate-based scRNA-sequencing of EpCAM^+^ChAT^BAC^-eGFP^+^ tuft cells isolated from mouse small intestinal tissue. Event density is indicated by color gradient. Sorted events are denoted in black. b. Mean fluorescence signal (eGFP) of cells from indicated UMAP clusters measured by FACS prior to plate-based scRNA-sequencing (ns not significant, **** p<0.001, t-test). c. Box plot indicating the scores by which various small intestinal cell type signature^14^ can be detected in single cells of the indicated tuft d. clusters as identified in the ChAT^BAC^-eGFP dataset. e. Violin plots indicating the scores by which the zonated expression signatures along the crypt-villus axes^15^ are detected in single cells of the three tuft clusters found in the ChAT^BAC^-eGFP dataset.

**Extended Data Figure 2.**
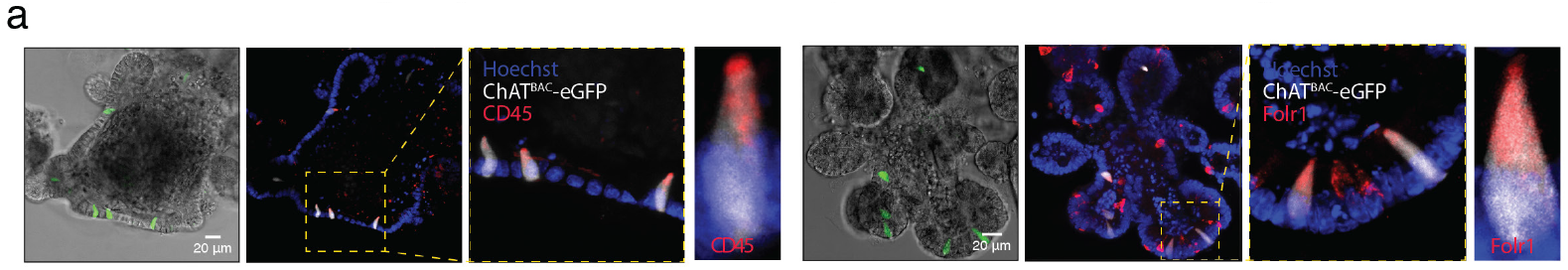
Mature tuft cell markers are expressed in mouse small intestinal organoids Related to Figure 3. a. Microscopy pictures of mouse small intestinal organoids from ChAT^BAC^-eGFP mice showing brightfield image (grey) overlayed with fluorescent GFP signal (green). From each organoid the corresponding fluorescent images are shown in three magnifications (GFP signal in white) with immunofluorescence staining against indicated tuft cell markers CD45 or FOLR1 (red). For counterstaining, nuclei (blue) are labelled with Hoechst 3342.

**Extended Data Figure 3.**
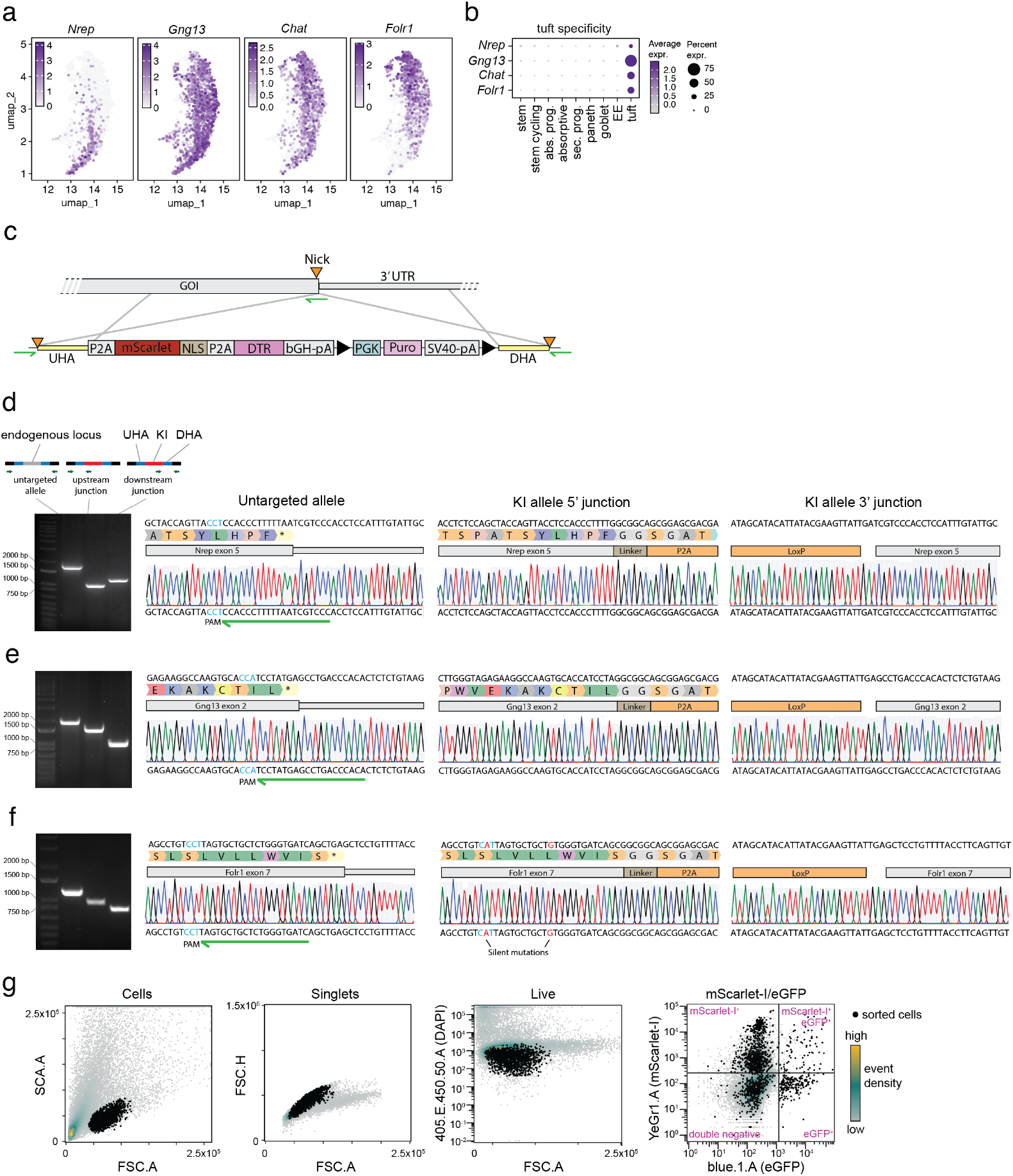
Tuft type-specific reporter knock-ins in mouse small intestinal organoids Related to Figure 3. a. UMAP of integrated *in vivo* tuft cell cluster (Figure 1g), overlayed with heatmap showing relative expression of selected reporter genes for prospective tuft type-specific knock-ins (Figure 1g). b. Relative expression level of selected tuft type-specific reporter genes in various intestinal cell types featured in the integrated *in vivo* mouse small intestine single-cell transcriptome dataset (Figure 1g). Average expression is represented by dot color. Percentage of expressing cells is denoted by dot size. EE: enteroendocrine, abs. prog.: absorptive progenitor, sec.prog.: secretory progenitor. c. Schematic representation of genetic knock-in strategy using in-trans paired nicking, where Cas9D10A nickase generates single strand breaks (‘nick’, orange triangles) in genomic target site, as well as at the extremities of the homology arms in the knock-in donor template. Guide RNA (green arrows) targets vicinity of C-terminal STOP-codon. Black triangles indicate LoxP sites. UTR: untranslated region, UHA: upstream homology arm, DHA: downstream homology arm, NLS: nuclear localization signal, DTR: diphtheria toxin receptor, bGH-pA: polyadenylation signal of bGH, PGK: PGK promotor, Puro: puromycin resistance cassette and SV40-pA: polyadenylation signal of SV40. d. Validation of correct genomic integration of heterozygous knock-in in Nrep by PCR (left) and Sanger sequencing of the corresponding PCR amplicons. Untargeted allele remains unaffected with in-trans paired nicking. UHA: upstream homology arm, DHA: downstream homology arm, KI: integrated knock-in, PAM: protospacer adjacent motif. Guide RNAs are indicated with green arrows. e. As in d, but for Gng13. f. As in d, but for FolrI. g. Gating strategy used during FACS prior to plate-based scRNA-sequencing of fluorescent cell populations from tuft type-specific reporter organoids. Event density is indicated by color gradient. Sorted events are denoted in black. Plot includes subsampling of events from all three organoid reporter lines and both medium conditions.

**Extended Data Figure 4.**
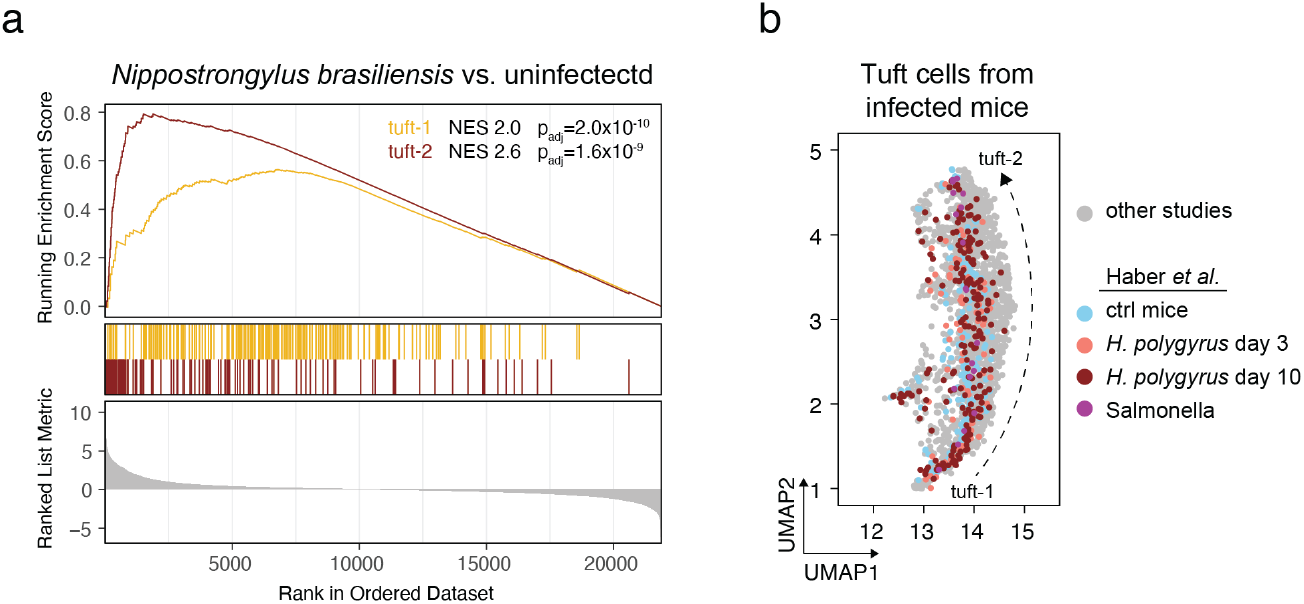
Type 2 cytokine-driven tuft cell hyperplasia leads to increased numbers of both tuft-1 and tuft-2 cells *in vivo* Related to Figure 4. a. Gene set enrichment analysis of tuft-1 and tuft-2 signatures^14^ in mice infected with *Nippostrongylus Brasiliensis*. Bulk RNA-sequencing data of infected and control mice was obtained from E-MTAB-9183^23^. b. Single-cell RNA sequencing profiles of tuft cells isolated from mice infected with *Heligmosomoides polygyrus* (obtained from GSE92332^14^) highlighted in UMAP of tuft cell cluster from integrated mouse *in vivo* single-cell transcriptome dataset (Figure 1g).

**Extended Data Figure 5.**
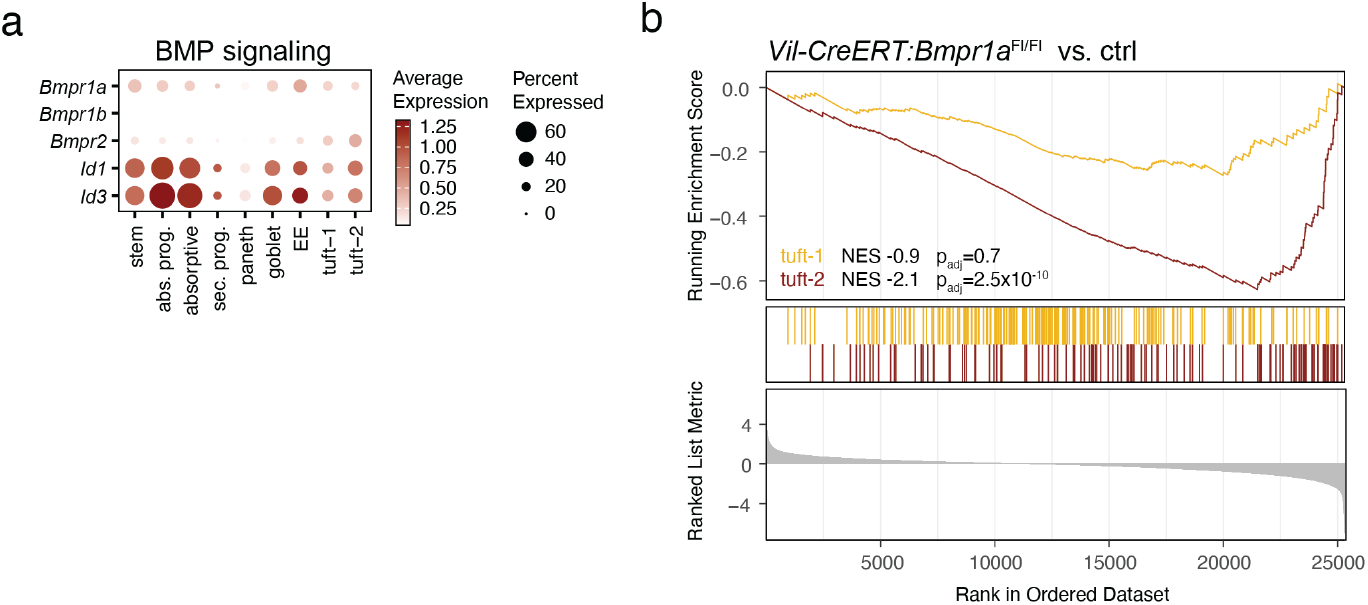
Tuft cell maturation to tuft-2 phenotypes depends on BMP signaling Related to Figure 5. a. Expression level of BMP receptors and BMP responsive genes in indicated cell types extracted from the integrated *in vivo* mouse small intestine single-cell transcriptome dataset (Figure 1g). Average expression is represented by dot color. Percentage of expressing cells is denoted by dot size. b. Gene set enrichment analysis of tuft-1 and tuft-2 signatures^14^ in bulk RNAseq dataset of intestinal epithelium depleted for *Bmpr1a* (*Vil-CreERT:Bmpr1a*^FL/FL^) vs. control mice^25^. For current analysis, RNA data from multiple diets were included in the comparison between *Bmpr1a* knockout and control groups. Upon perturbed BMP signaling, tuft-2 signature genes are the most downregulated.

**Extended Data Figure 6.**
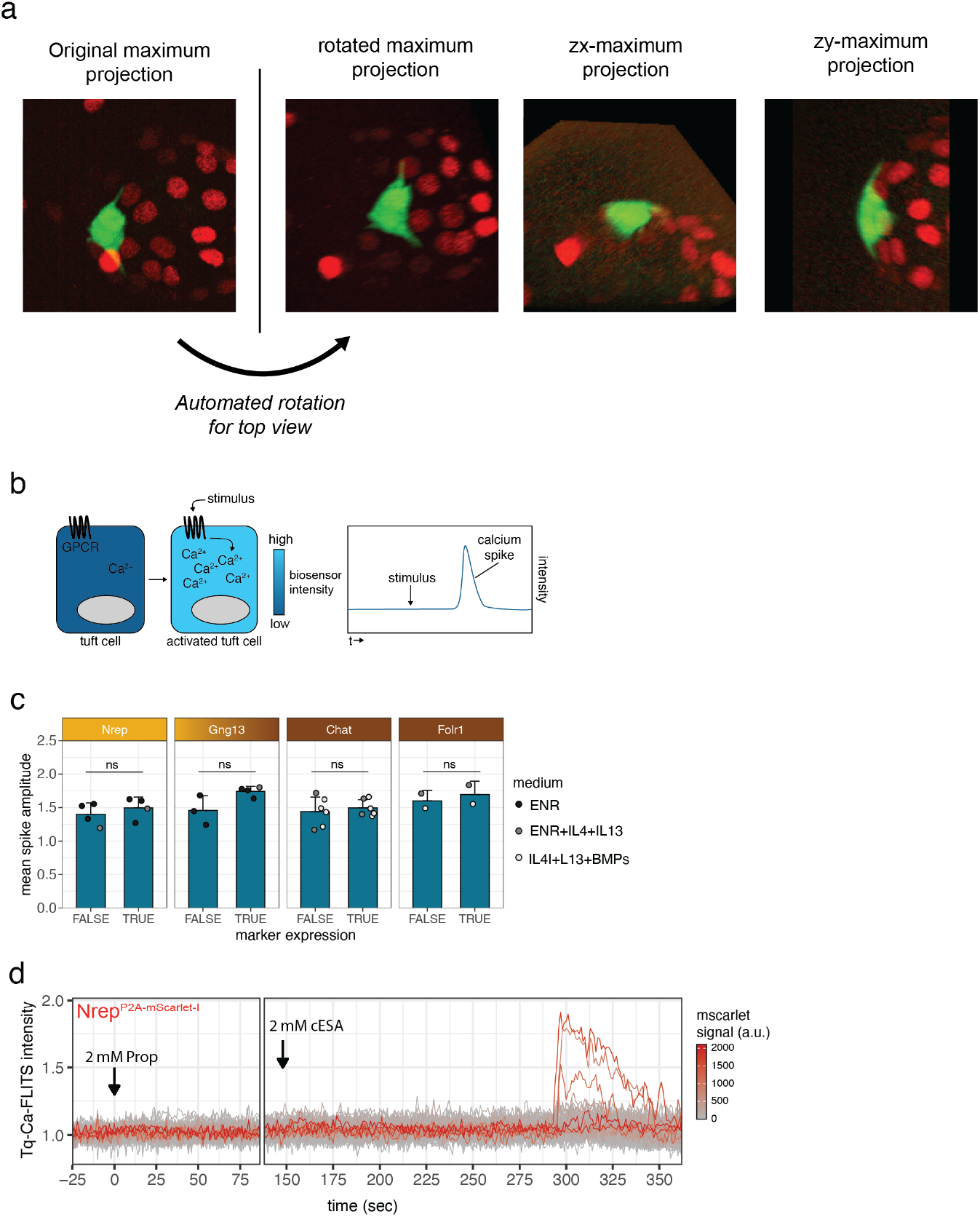
Dynamic tuft cell protrusion analysis and calcium signaling dynamics in organoid monolayers following stimulation with cESA and propionate Related to Figure 6. a. Stills examplifying automated rotation algorithm used to visualize the position of tuft cell protrusions within the organoid epithelial layer. Protrusions wrap around the basal side of the organoid. Green: ChAT^BAC^-eGFP, Red: H2B-mScarlet-I. b. Schematic of tuft cell activation measurements with Tq-Ca-FLITS calcium biosensor. c. Bar diagram depicting mean amplitude of calcium spike following stimulation with 1.5 mM cESA. Per bar, each point represents a separate experiment and is colored per medium used. Data are represented as mean ± SD (ns not significant, t-test). d. Single-cell Tq-Ca-FLITS intensity traces following sequential stimulation with 2 mM propionate, followed by 2 mM cESA. Traces are colored by mScarlet-I signal measured in the corresponding cell.

